# Localization of candidate clock neurons in Antarctic krill (*Euphausia superba*)

**DOI:** 10.64898/2026.02.26.708226

**Authors:** Lukas Hüppe, Nils Reinhard, Annika Karl, Valentina Kirsch, Laura Wollny, Amy Palmer, Dirk Rieger, Pingkalai R. Senthilan, Charlotte Helfrich-Förster

## Abstract

Organisms are exposed to predictable daily and seasonal environmental oscillations. Biological clocks enable organisms to anticipate these changes and coordinate physiology and behaviour accordingly. While circadian mechanisms are well studied in terrestrial model organisms, little is known about the neuronal organisation of biological clocks in non-model species, especially those inhabiting marine environments. Antarctic krill (*Euphausia superba*) is central to the functioning of the Southern Ocean ecosystem and relies on precise timing to cope with the extreme, high-latitude fluctuations in photoperiod, food availability, and sea-ice cover in its habitat. Despite evidence for circadian and seasonal rhythms in krill behaviour and physiology, the neuronal architecture underlying these timing processes has remained unresolved. In this study, we use *in situ* hybridisation and antibody staining to characterise the circadian clock in the krill brain. Immunostaining with an antibody against crustacean β-pigment-dispersing hormone (β-PDH) reveals distinct clusters of PDH-positive neurons in the eyestalks and dorsal central brain, along with an extensive PDH-positive fibre network. We further localise transcripts of the core clock genes *cryptochrome-2* (*cry2*) and *period* (*per*) in cell clusters in the eyestalks, which also include the PDH-positive neurons. More specifically, PDH-positive neurons are a subgroup of the *cry2*-and *per*-positive cells. Together, these findings provide the first description of the neuronal architecture of the circadian clock in Antarctic krill and establish essential groundwork for future studies on biological timing and environmental adaptation.

## Introduction

Throughout evolution, organisms are subjected to regular environmental oscillations at daily and seasonal time scales, like changes in light intensity, temperature, humidity, and food availability. The underlying cause for these oscillations is the rotation of the Sun-Earth-Moon system, which makes these oscillations highly predictable and led to the evolution of biological clocks (Patke et al., 2020). Biological clocks are present across the tree of life and share a common structure: A self-sustaining endogenous oscillator that can be synchronized by external cues (*Zeitgebers*) generates rhythms in, e.g., gene expression or neuronal activity. These rhythms coordinate the organism’s physiology and behaviour with the appropriate phase of daily or seasonal cycles (Dunlap et al., 2004). Accordingly, endogenous clocks allow an organism to predict, rather than simply react to, regular environmental changes, enhancing an organism’s ability to adapt to its environment, hence increasing its fitness (Emerson et al., 2008; Horn et al., 2019; Jabbur et al., 2024).

By far the best studied timing mechanism is the circadian clock, which generates rhythms with a period length of ∼24 h and synchronizes to the daily light-dark cycle. Today, the circadian clock mechanism is known in great detail for a few terrestrial model species, such as *Drosophila melanogaster*, *Mus musculus*, *Neurospora crassa*, or *Arabidopsis thaliana*, which have been chosen for their genetic tractability, easy husbandry, and experimentally accessible cellular or neural systems (Johnson et al., 2026). These models have led to a deep understanding of circadian clocks (Beer & Helfrich-Förster, 2020).

Accelerating anthropogenic pressures, such as rapid global warming, challenge organisms. Their adaptive capacity is a major determinant of species resilience to these changes. As a result, there is growing momentum to extend chronobiological research beyond traditional model organisms and to species across diverse taxa and ecosystems (Häfker et al., 2022; Häfker & Tessmar-Raible, 2020; Helm et al., 2013; Helm & Liedvogel, 2024; Helm et al., 2017; Johnson et al., 2026; Kronfeld-Schor et al., 2017).

Antarctic krill (*Euphausia superba*, hereafter “krill”) is an up to 6 cm long marine, pelagic crustacean that belongs to the Euphausiacea, an order within the Malacostraca (Siegel, 2016). It is endemic to the Southern Ocean, a high-latitude habitat with strong seasonal fluctuations in daylength (photoperiod), food availability, and sea-ice cover. Krill efficiently makes use of the high primary productivity in the summer months and serves as an important prey for apex predators, including whales, penguins, seals, and seabirds, thus forming a crucial link between low trophic levels and higher consumers (Ballerini et al., 2014; Murphy et al., 2007). Krill thrives in the harsh conditions of the Southern Ocean, and with estimates of >300 Mio tons, their biomass is considered the highest of any wild living single species on Earth (Atkinson et al., 2009; Bar-On et al., 2018). Its close, rhythmic adaptation to the daily and seasonal fluctuations of the environment is thought to be key to krill’s ability to sustain this biomass in such a challenging habitat (Ducklow et al., 2007). Rhythmic regulation of physiology and behaviour of krill has been observed at both daily and seasonal time scales. For example, every night, large krill swarms migrate tens to hundreds of meters to the upper parts of the water column to feed on phytoplankton and micro zooplankton. With sunrise, krill return to deeper parts to hide in the dark from visually hunting predators (Bahlburg et al., 2023; Cisewski et al., 2010; Everson, 1983). Due to their large biomass, this synchronized behaviour, known as diel vertical migration (DVM), has major impacts on predator-prey interactions (Annasawmy et al., 2023; Cox et al., 2009; Nichols et al., 2022), the recycling of nutrients, and the efficiency of carbon export (reviewed in Cavan et al., 2019). Seasonally, krill shows a pronounced regulation of physiological functions, including halted development, reduced metabolism, and altered lipid dynamics, which are closely aligned with the seasonal availability of food (Kawaguchi et al., 2007; Meyer et al., 2010). Most importantly, the circadian clock has been shown to drive daily rhythms in swimming behaviour, related to DVM (Hüppe et al., 2025; Piccolin et al., 2020) and suggested to be involved in the photoperiodic control of seasonal metabolism and reproduction (Höring et al., 2018; Piccolin, Suberg, et al., 2018; Teschke et al., 2007, 2008). The molecular mechanism of circadian clocks is based on transcriptional-translational feedback loops (TTFL), in which complex interactions of clock genes and their protein products result in self-sustained ∼24h rhythms. In animals, TTFLs generally comprise positive elements that promote transcription and negative elements that inhibit it. This mechanism is largely conserved across taxa; however, systematic differences exist, particularly with respect to the negative elements. In vertebrates and most arthropods, Period (PER) and the light-insensitive mammalian Cryptochrome (mCRY or CRY2) act as transcriptional repressors, while Clock (CLK) and Basic helix-loop-helix arnt-like protein/Cycle (BMAL/CYC) act as transcriptional activators (Kotwica-Rolinska, Chodakova, et al., 2022; Patke et al., 2020). Similarly, this is also the case in various Crustacea (Christie et al., 2013; Christie, Yu, Pascual, et al., 2018; Hunt et al., 2019; Nesbit & Christie, 2014; O’Grady et al., 2016; Sbragaglia et al., 2015; Tilden et al., 2011), including Euphausiacea (Biscontin et al., 2017; Christie, Yu, & Pascual, 2018). In contrast, the well-known *Drosophila* clock mechanism represents a special case where only the light-sensitive *Drosophila-*type Cryptochrome (dCRY or CRY1) is present, and which, by interaction with Timeless (TIM), resets the clock in a light-dependent manner (Emery et al., 1998). Interestingly, krill as well as several other insect and crustacean species, possess both forms of Cryptochromes, representing an evolutionarily ancient molecular clock (Biscontin et al., 2017; Deppisch et al., 2022; Mazzotta et al., 2010; Yuan et al., 2007). In krill, CRY2 in combination with PER is thought to be the major repressor of CLK-BMAL/CYC activity, while binding of TIM is thought to stabilize the complex (Biscontin et al., 2017). This is similar to the mechanism described for the monarch butterfly *Danaus plexippus* (Brady et al., 2021; Zhu et al., 2008) and the marine isopod *Eurydice pulchra* (Zhang et al., 2013).

The molecular clock ticks in many tissues and cells; however, only a small number of neurons in the brain control rhythmic behaviour and coordinate the daily oscillations in other tissues (Albrecht, 2012; Tomioka et al., 2012). As known from insect studies, these clock neurons are generally organized into distinct clusters, typically found associated with the optic lobes (equivalent to the eyestalks of Crustacea) and protocerebrum (reviewed in Helfrich-Förster et al., 1998; Numata et al., 2015). The single neurons are strongly interconnected through, e.g., synaptic and paracrine signalling within a clock network. This whole network then generates coherent rhythmicity and drives the rhythmic biology of the organism (reviewed in Helfrich-Förster & Reinhard, 2025). The best studied neuropeptide of the circadian clock is pigment-dispersing factor (PDF), which today is known to be involved in phasing and synchronizing the clock neurons and the control of rhythmic behaviour (Liang et al., 2016; Renn et al., 1999; Rojas et al., 2019; Stengl & Arendt, 2016; Vaze & Helfrich-Forster, 2021). Interestingly, modulations of the PDF neuronal network have been implicated in the ability of high latitude flies to synchronize their locomotor activity to long photoperiods (Menegazzi et al., 2017). Further, PDF has been shown to provide a link between the circadian clock and the timing of seasonal events (i.e., photoperiodism; Hasebe et al., 2022; Hidalgo et al., 2023; Kotwica-Rolinska, Damulewicz, et al., 2022; Nagy et al., 2019). PDF-like peptides have first been isolated from the eyestalks of crustaceans, and one form, termed β-pigment-dispersing hormone (β-PDH), has been identified as the most common one, well-known for its effect on the dispersion of retinal and chromatophore pigments in crustaceans (Rao & Riehm, 1993). Besides their structural similarity, functional similarity has been suggested between crustacean PDH and insect PDF. For example, head extracts from several insects have been shown to be able to induce pigment dispersion in crustaceans (reviewed in Rao, 2001), while β-PDH from the intertidal crab *Cancer productus* is able to restore rhythmic activity in *Drosophila melanogaster Pdf*-null mutants (Beckwith et al., 2011). Also in krill, a β-PDH sequence has been identified, but its spatial expression and function remain unknown (Hunt et al., 2017; Toullec et al., 2013).

Most knowledge of circadian clock neuronal networks comes from insect studies, while much less is known about these systems in malacostracan crustaceans. Given their shared pancrustacean ancestry and conserved brain architecture (e.g., reviewed in Harzsch & Krieger, 2021; Strausfeld, 2009; Strausfeld, 2012), however, key features of the circadian clock system are likely conserved between the two groups. This has been proposed particularly for the network of PDH-expressing neurons (Harzsch et al., 2009), which has been described in various malacostracan crustaceans, including *Carcinus maenas* (Alexander et al., 2020; Mangerich & Keller, 1988; Mangerich et al., 1987), *Cancer productus* (Hsu et al., 2008), *Orconectes limosus* (de Kleijn et al., 1993; Mangerich & Keller, 1988; Mangerich et al., 1987), *Homarus americanus* (Harzsch et al., 2009), *Cherax destructor, Procambarus clarkii* (Sullivan et al., 2009), and *Procambarus virginalis* (Luna et al., 2010). Across Malacostraca, the general brain ground plan is highly conserved (Schmidt, 2016), yet the relative volume of specific sensory neuropils varies in correspondence with ecological specialization (Chittka & Niven, 2009; Niven & Laughlin, 2008; Sandeman et al., 2014), as demonstrated across shrimp-like crustaceans occupying a range of niches (Krieger et al., 2020; Machon et al., 2019; Meth et al., 2017). While only little is known about the neurobiology of *E. superba*, its brain appears to follow the malacostracan ground plan (Sandeman et al., 1993). The distinct pelagic niche of euphausiids likely entails differences in the relative development of certain sensory centres — such as a disproportionate investment in visual processing areas, given the exceptional importance of light as an environmental cue in open-ocean habitats. As the relevant timing processes are orchestrated at the level of individual neurons, establishing a foundational understanding of its neuroanatomy and organization of its neuronal circadian system is a critical step for any deeper mechanistic insight.

In this paper, for the first time, we revealed the spatial organization of putative circadian clock neurons in the brain of *E. superba*. First, we used immunohistochemistry and *pdh in situ* hybridization to reveal distinct clusters of PDH-positive neurons in the eyestalks and the dorsal central brain, as well as an extensive fibre network. Subsequently, we localize the transcripts of the core clock components *cry2* and *per* and show that they partially co-localize with PDH-positive neurons. This study reveals the neuronal architecture of the krill circadian clock and thereby provides a fundamental basis for future studies of biological timing in krill.

## Results

Antarctic krill has a shrimp-like habitus, which is divided into the cephalothorax and the pleon with six segments (Figure 1a). The Cephalothorax bears eight pairs of thoracic legs, the endopodites of which six are transformed to form a feeding basket, used to filter the water for phytoplankton and microzooplankton. The pleon bears five pairs of swimming legs (pleopods), which are the main means of propulsion (Figure 1a, white arrows). The dissected central nervous system reveals the brain, two circumoesophageal (blank arrowheads), the suboesophageal, eight thoracic (T1-T8), and six pleonic (PL1-PL5 + terminal pleonic ganglion, TPG) ganglia along the ventral nerve cord (VNC; Figure 1b and c). The brain is dominated by the large, stalked compound eyes, whose volume well exceeds that of the central brain, as well as by the nerves of the antenna 1 (ANT1, Figure 1a to c), which exit the central brain anteriorly, and the antenna 2, which exit the posterior brain ventrally (ANT2; Figure 1b).

**Figure 1:**
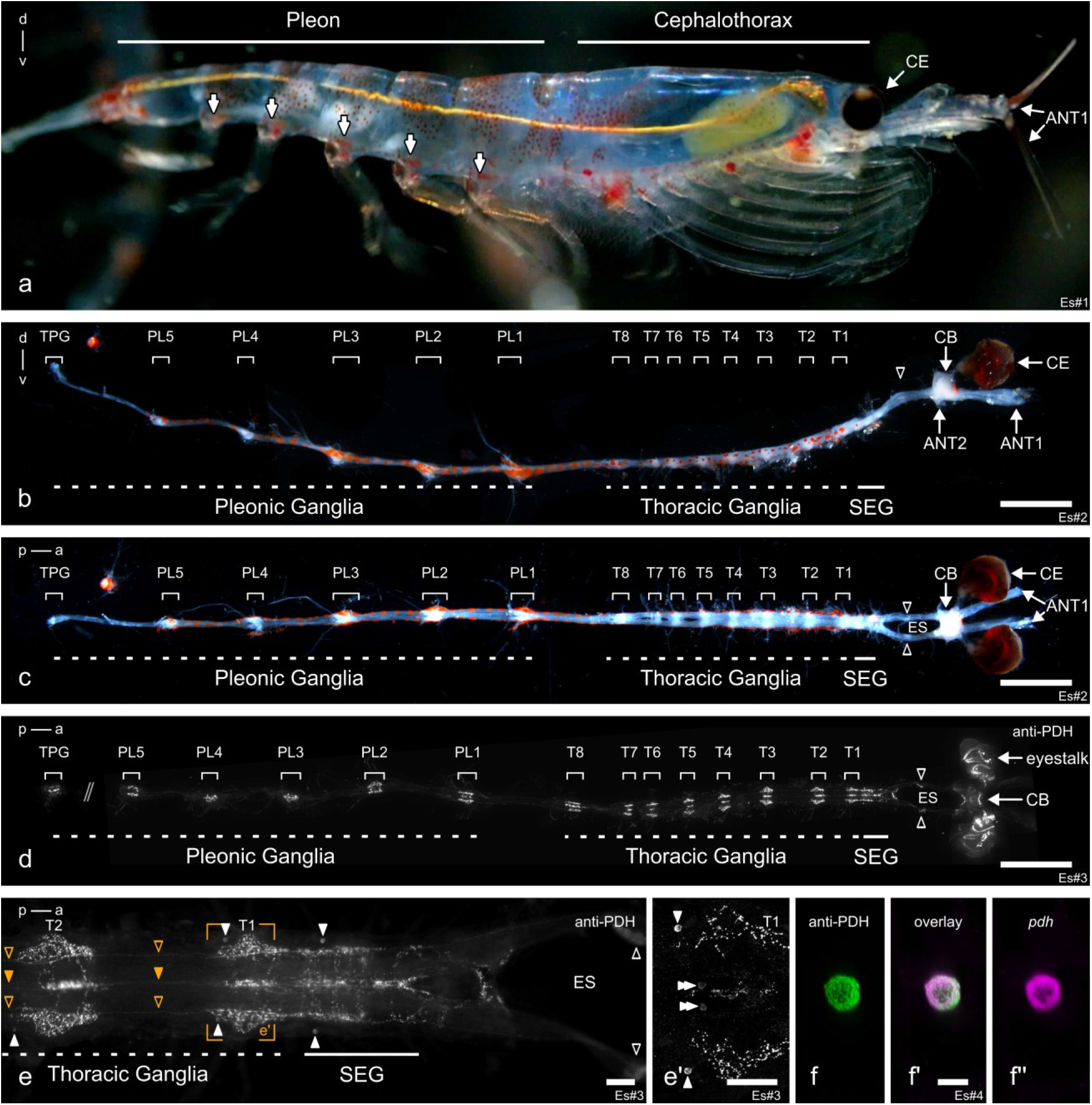
Gross morphology of *Euphausia superba* and its central nervous system. **a)** Image of a swimming *E. superba*, facing with dorsal to the top and anterior to the right. Thick white arrows point to the five pairs of swimming legs (pleopods) of the pleon. **b, c)** Lateral (b) and dorsal (c) view of the dissected central nervous system (CNS) of krill. The ventral nerve cord can be divided into suboesophageal (SEG), thoracic (T1-8), and pleonic (PL1-PL5 + TPG) ganglia. Arrows point to the compound eyes (CE), central brain (CB), and the 1^st^ and the base of the 2^nd^ antennae (ANT1 and ANT2, respectively). Blank arrowheads label the circumoesophageal ganglia in figure panels (b-e). Note the orange erythrophores distributed along the ventral nerve cord. **d)** Pigment-dispersing hormone-immunoreactivity (PDH-ir) within the CNS of krill. Note that the terminal pleonic ganglion (TPG) was recorded independently of the rest of the CNS due to restrictions in dissections. **e)** Detailed view of the SEG and the first two thoracic ganglia (T1, T2). PDH-ir cells are labelled by white arrowheads. Fibers connecting the ganglia in the lateral connectives are labelled by blank orange arrowheads, and fibres extending through the median neurite bundle by a filled orange arrowhead. The orange frame indicates the location of the magnified view in (e’). **e’)** shows a magnified view of T1 recorded with higher resolution. The two laterally located, strongly PDH-ir cells are labelled with arrowheads, and the two weakly PDH-ir cells close to the median bundle are labelled with double arrowheads. **f)** Representative cell (from T2) of one of the paired cells in the ventral nerve cord ganglia labelled by anti-PDH (green) and *pdh* in situ hybridization (magenta). The scalebar represents 3 mm for (b-d), 200 µm for (e and e’), and 50 µm for (f). Abbreviations: d, dorsal; v, ventral; a, anterior; p, posterior, ES, oesophageal foramen. Figure panels showing the same individual are indicated with the same Es#ID.

### An antibody against crab β-PDH reveals an extensive network of PDH-positive neurons throughout the krill central nervous system

To visualize potential clock neurons, we stained against the clock-related neuropeptide ß-PDH, as previous studies have shown strong interspecific cross-reactivity and even a functional conservation between crabs and flies (Beckwith et al., 2011; Dircksen et al., 1987; Harzsch et al., 2009; Helfrich-Förster, 1995; Hsu et al., 2008; Mangerich et al., 1987; Nussbaum & Dircksen, 1995). Immunostainings of the dissected central nervous system with an antibody against crab β-PDH (Wilcockson et al., 2011) show extensive PDH-immunoreactivity (PDH-ir) (Figure 1d). Dense innervation is found for the brain (Figure 1d and Figure 2) and both circumoesophageal ganglia (Figure 1d, e, blank arrowheads). Similarly, PDH-ir fibres strongly innervate each ganglion along the VNC in both lateral neuropils, the anterior and posterior commissures, and along the midline (Figure 1d, e). We found PDH-ir fibres extending in the median neurite bundle (Figure 1e, orange arrowhead) and the two lateral connectives (Figure 1e, blank orange arrowheads), linking all single ganglia along the VNC. We also identified a pair of strongly PDH-ir cells located laterally in the posterior region of all ganglia of the VNC, commonly positioned medially along the dorsal-ventral axis (Figure 1e, e’ and f, white arrowheads). Additionally, we found further ventrally a pair of weakly PDH-ir cells close to the median bundle (Figure 1e’ double arrow heads). We could clearly identify these weakly stained cells close to the median bundle in the SEG and the thoracic ganglia, however, we could not unambiguously identify the presence of these cells in the pleonic ganglia. As we only had few samples for the ventral nerve cord (N=3) and the cells are only visible at high resolutions, further studies are necessary to investigate the presence of PDH-ir cells close to the median bundle in the pleonic ganglia. While the strongly and weakly stained cells appear to contribute to the dense fibre networks in the ganglia, they do not seem to be their only source.

**Figure 2:**
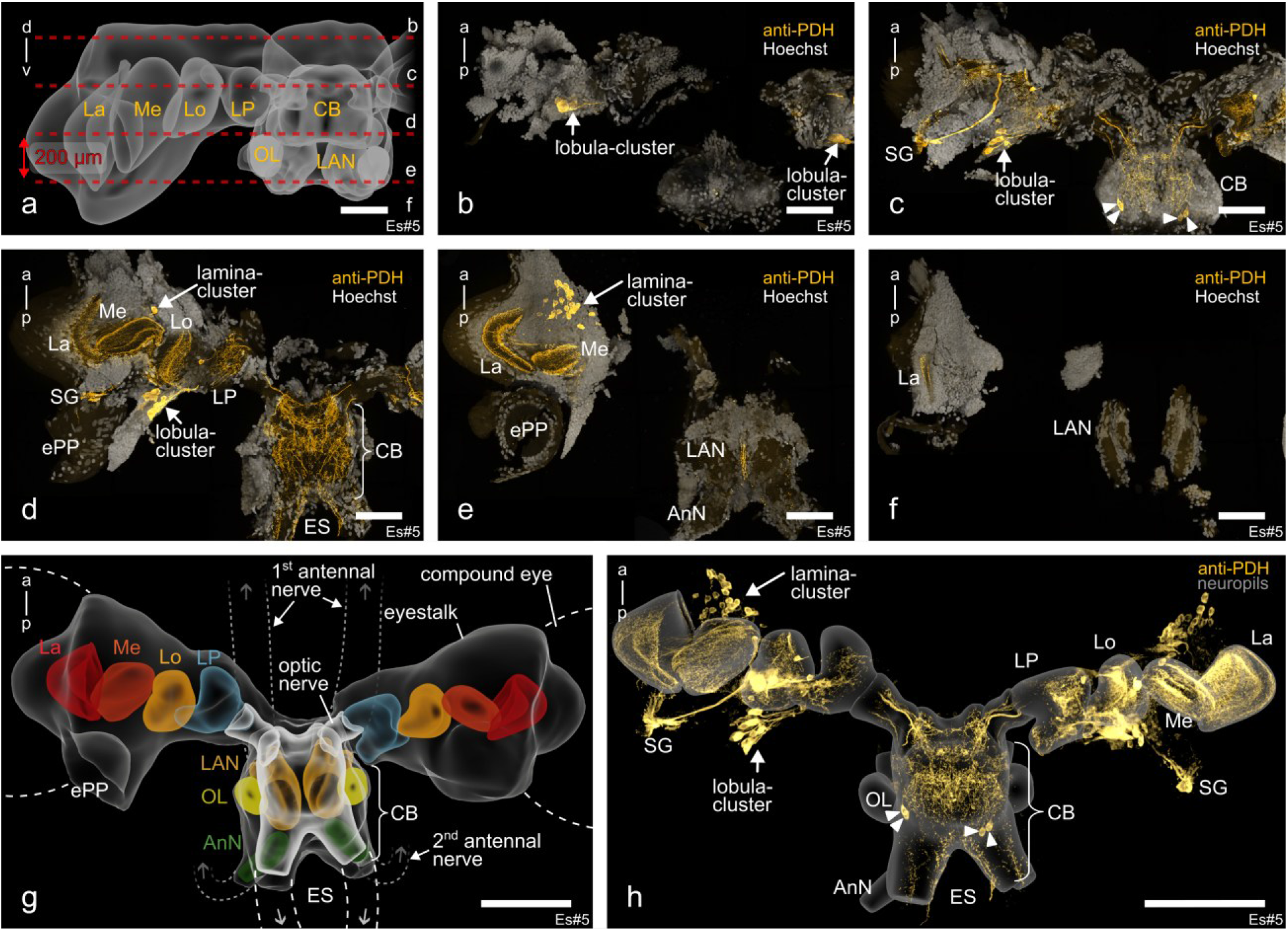
Overview of the PDH-network in the brain of krill. **a)** Frontal view of one eyestalk and the central brain (CB) showing the position of the five horizontal 200 µm sections depicted in (b-f). **b-f)** Horizontal vibratome sections of the brain (left eyestalk and central brain) stained against β-PDH (orange) with a nuclear counterstaining (Hoechst, grey). The sections show extensive PDH-immunoreactivity from the dorsal (b) to ventral (f) brain. In the eyestalk, two prominent cell clusters (lamina-cluster, lobula-cluster) can be found. **g)** Reconstruction of major neuropils found in the brain. The eyestalks contain, from lateral to medial, the lamina (La, red), medulla (Me, dark orange), the lobula (Lo, light orange), and the lateral protocerebrum (LP, light blue). The olfactory neuropils comprise the lateral antenna 1 neuropil (LAN, ochre), the olfactory lobes (OL, yellow), and the antenna 2 neuropil (AnN, green). The colour scheme is based on Kenning et al. (2013). Grey arrows indicate continuation of cut connectives/ nerves. Note that, due to the eyestalks’ fragile connection to the central brain, the right eyestalk was rotated horizontally approximately 40° counterclockwise during processing. **h)** 3D projection of the whole PDH-network spanning the krill brain. The neuropils from (g) are shown in grey for orientation. Scale bars in (a-f) represent 250 µm and 500 µm in (g) and (h). Abbreviations: ES, oesophageal foramen; SG, sinus gland; ePP, eyestalk photophore. Figure panels showing the same individual are indicated with the same Es#ID.

The most elaborate arborization pattern could be found within the brain. Widespread PDH-ir fibres innervated the central brain and the eyestalks of krill (Figure 1d and Figure 2). Since the brain was too thick for whole-mount imaging, we decided to use 200 µm horizontal vibratome sections (Figure 2a) to investigate further the arborisations of the PDH-network within the central brain and the eyestalks (Figure 2b-f, h). Based on nuclear (Hoechst) and neuropil staining (anti-synapsin; Klagges et al., 1996) combined with autofluorescence, we could identify major brain regions, including the visual neuropils and neuropils of the lateral protocerebrum (Figure 2a, g, Figure 2-figure supplement 1a-e). As with most other malacostracan crustaceans, the krill brain can be divided into three regions: the protocerebrum, which is further subdivided into the lateral protocerebrum (comprising the medulla terminalis and the mushroom body) and the medial protocerebrum (which constitutes the anterior part of the central brain), the deutocerebrum (which hosts the lateral antenna 1 neuropils and olfactory lobes), and the tritocerebrum (which constitutes the posterior central brain, the antenna 2 neuropils and the oesophageal connectives). PDH-ir fibres in the eyestalks clearly outline the visual neuropils, namely lamina, medulla, and lobula (Figure 2b-f, h).

At the base of the eyestalk, the lateral protocerebrum is found (Figure 2g, Figure 2-figure supplement 1d) which appears strongly innervated by PDH-ir fibres (Figure 2d, h). The fibres originate from two major cell clusters within the eyestalk: one cluster is located anteriorly between medulla and lamina and closely associated with the lamina, hence called the lamina-cluster, and the other is closely associated with the lobula, hence called the lobula-cluster. The following paragraph provides a detailed description of the clusters and their arborisations. To verify that the PDH-ir cells are indeed PDH-positive, we performed *pdh* RNA *in situ* hybridization on horizontal vibratome sections of krill brains (Figure 2-figure supplement 2). All strongly PDH-ir cells were also labelled by *pdh in situ* hybridization, independent of the cluster (Figure 1f and Figure 2-figure supplement 2a-j). Our data indicate that the cells that are strongly labelled by anti-β-PDH are indeed PDH-positive. However, we found several weakly PDH-ir cells in the central brain and the lobula-cluster, where we could not clearly determine the co-localization of anti-PDH and *pdh in situ* labelling (Figure 2-figure supplement 2g’, i’ and j’; blank arrowheads). While there is the possibility that these neurons do not express *pdh* and are only labelled due to cross reactivity of the anti-β-PDH antibody with other peptides or proteins, it might well be that the expression is too low for us to detect via RNA *in situ* hybridization. The latter seems more likely since all strongly labelled cells were always co-localizing with *pdh* and could be reliably detected. The weakly stained neurons, on the other hand, were more variable and could not always be detected depending on the background fluorescence, settings and method used for acquisition. We therefore focus in the following on the cells that show both strong PDH-ir and express *pdh*.

### The krill eyestalks contain the vast majority of PDH-positive neurons

The lamina-cluster is located in the ventral anterior eyestalk and is the largest and most prominent of the PDH-positive cell clusters in the krill eyestalks (Figure 2d, e, h, Figure 3a-d, and Figure 2-figure supplement 3 and 4). We could reliably detect between 46 to 67 homogenously stained cells per hemisphere (M = 59.0 n = 6) with a cell body diameter ranging between 25.8 µm and 57.3 µm (M = 38.6 µm, mean SD = 6.3 µm) (Figure 3-figure supplement 1 and 2). While the cells of this cluster appear to be organized in distinct groups (Figure 3a, b) and likely comprise multiple subclusters, we currently lack robust characteristics to reliably separate them; it is therefore described here as a single cluster. At the proximal side of the lamina, we found several PDH-positive cells of this cluster distributed along the surface (filled arrowheads in Figure 3a-c). From the lamina, the cluster extends medially to the antero-ventral base of the medulla (Figure 3a-c). The projections of parts of the lamina-cluster neurons appear to extend along the surface of the medulla before they innervate the lamina (Figure 3d). Some of these projections innervate the medulla, next to the lamina (blank arrowheads in Figure 3c). The PDH-positive fibres that extend between the lamina and the medulla cross within the outer and inner visual chiasm, respectively (single and double asterisks in Figure 3a, respectively).

**Figure 3.**
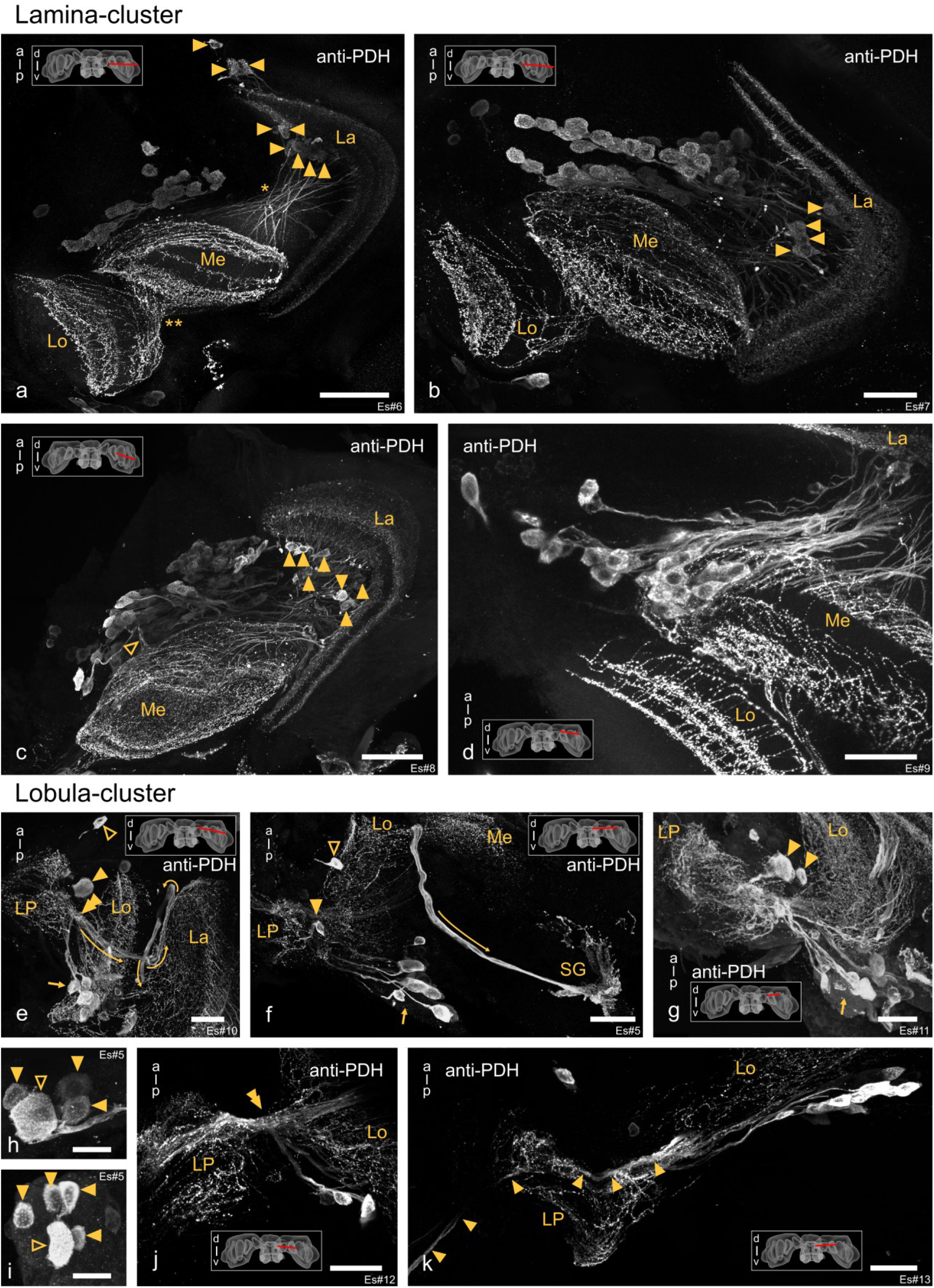
Detailed morphology of PDH-positive cell clusters and their projections in the krill eyestalks. a-d) Maximum intensity projections showing PDH-positive cells and their projections in the lamina-cluster. **a-c)** Filled arrowheads point to cells distributed along the proximal surface of the lamina, open arrowhead in (c) points to projections into the medulla. Single and double asterisks in (a) indicate the outer and inner visual chiasm, respectively. **e-k)** Maximum intensity projections of PDH-positive cells and their projections in the lobula-cluster. **e-g)** single arrows point to lobula-subcluster 1 at the posterior side of the eyestalk and filled arrowheads point to cells of lobula-subcluster 2, projecting into the lateral protocerebrum. Blank arrowheads in (e-f) point at single PDH-positive cells between the lobula and lateral protocerebrum. Double arrowheads in (e) and (j) point towards the dense fibre knot formed by projections of lobula-subcluster 1 cells, whereas yellow lines in (e-f) outline their projection lateral of the knot. **h, i)** Maximum intensity projection of subcluster 2 of the lobula-cluster showing five PDH-positive cells, including one characteristic large cell body (open arrowhead) and four smaller cells (filled arrowheads). **k)** Projections of lobula-subcluster 1 through the eyestalk towards the central brain (filled arrowheads). Scale bars represent 200 µm in (a-c), and 100 µm in (d-g, j, k). Scalebars in (h-i) represent 50 µm. Abbreviations: La, lamina; Me, medulla; Lo, lobula; LP, lateral protocerebrum; SG, sinus gland; a, anterior; p, posterior. Figure panels showing the same individual are indicated with the same Es#ID.

The lobula-cluster lies further dorsally in the eyestalk posterior to the lobula and can be divided into two distinct subclusters (Figure 3e-g). Subcluster 1 comprises with 21 to 56 cells per hemisphere (M = 34.8, n = 6) most of the cells close to the lobula (arrows in Figure 3e-g, Figure 3-figure supplement 1 and 2). In contrast, subcluster 2 is located further anterior in the dorsal eyestalk (filled arrowheads in Figure 3e-g) and comprises only four to seven PDH-positive cells per hemisphere (M = 5.8, n = 6). One of the cells of subcluster 2 is much larger than the others (blank arrowhead in Figure 3h and i). Compared to the lamina-cluster most cells of the lobula-clusters are overall larger, but they vary strongly in size and staining intensity. Cells of subcluster 1 range between 24.3µm to 67.2µm in diameter (M = 44.0 µm, mean SD = 10.2 µm) and cells of subcluster 2 range between 35.7 µm to 74.2 µm (M = 48.2 µm, mean SD = 14.7 µm) in cell diameter (Figure 3-figure supplement 1 and 2). Subcluster 1 and subcluster 2 can be well distinguished in datasets in which cells of subcluster 2 are densely grouped together or the entire brain with the eyestalks was reconstructed from the single slices. However, this appeared more difficult when the cells of subcluster 2 were more dispersed, or no reconstruction of the entire brain was possible. Hence, we will refer to both subclusters together as loblua-cluster and only refer to them singularly when we could clearly distinguish between them.

Cells of the lobula-cluster (subcluster 1 and 2) send projections towards the lateral protocerebrum (Figure 3e-g). Between the lobula and lateral protocerebrum, these form a dense knot, from which a thick fibre bundle projects medially and laterally (double arrowheads in Figure 3e and j). Lateral to the knot, the fibre bundle passes the lobula, turns anterior before it loops back posterior between the lobula and medulla, targeting the sinus gland, which is situated between the lamina and the eye photophore in the posterior eyestalk (Figure 2d; lines in Figure 3e and f). Medially of the knot, the fibre bundle innervates the lateral protocerebrum (Figure 3e and j). Parts of this bundle appear to pass the lateral protocerebrum and extend through the eyestalk into the central brain (arrowheads in Figure 3k). However, due to the high number of fibres, the density of the fibre bundle and the dense arborisations in the lateral protocerebrum, we were unable to track the path of single fibres unambiguously.

Aside from the large cell clusters, two to six strongly PDH-positive single cells (M = 4.5, n = 6) were regularly found further apart of the defined clusters. Mostly they were located in the region between the lobula and lateral protocerebrum, and appeared to project towards the lateral protocerebrum (open arrowheads in Figure 3e and f, light-blue cells in Figure 3 figure supplement 1 and 2). Due to the variations in sectioning and their cell body position, we could not determine if they are indeed just single cells or part of the described clusters.

In contrast to the eyestalks, strong PDH-ir in the central brain is in most cases restricted to 2 cells per hemisphere (in some cases up to 4 cells) found dorsally in the medial protocerebrum (filled arrowheads in Figure 4a, 6i’ and j’). They project ventrally into the anterior protocerebrum (blank arrowheads in Figure 4a), where they cross fibres that extend from the eyestalks into the medial protocerebrum. Anterior to these fibres, the projections of the PDH-positive cells disappear in our preparations. The fibres from the eyestalks innervate a pair of granular, strongly PDH-ir structures, which lie dorsally in the medial protocerebrum (Figure 4a, asterisks). These structures strongly resemble the brain photoreceptor neuropil as described for the crayfish *Cherax destructor* and *Procambarus clarkii* (Sullivan et al., 2009).

**Figure 4:**
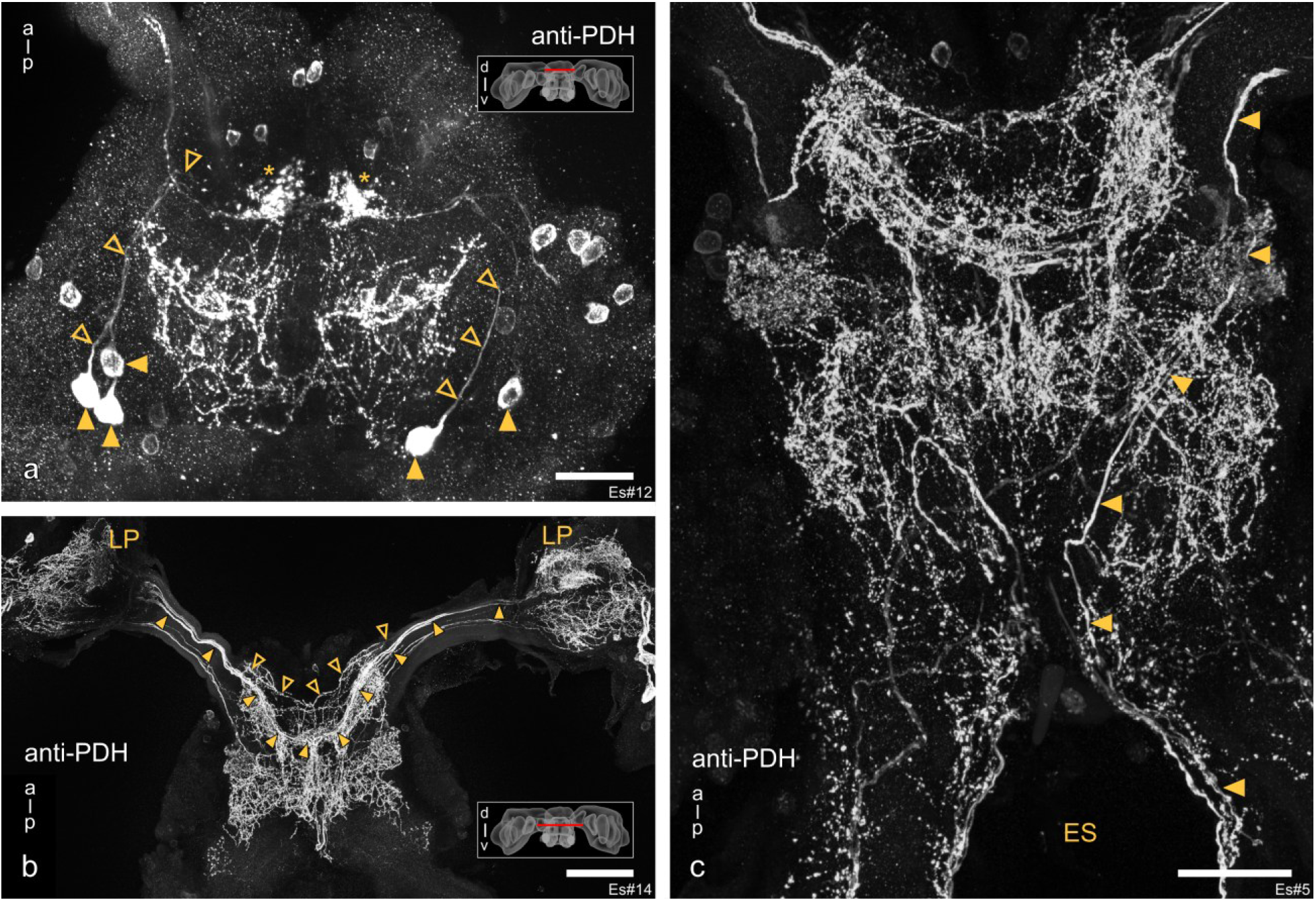
Detailed morphology of PDH-positive cells and projections in the krill brain a-c) Maximum intensity projections of the PDH-positive network in dorsal (a) and medial to ventral regions of the central brain (b-c). **a)** Filled arrowheads point to PDH-positive cells found dorsal in the medial protocerebrum, while blank arrowheads point to their projections. Asterisks indicate a paired, granular, PDH-ir structure which is targeted by fibres coming from the eyestalks. **b)** PDH-ir fibres originating from the lateral protocerebrum form dense arborisations in the medial protocerebrum. Filled and blank arrowheads indicate the paths of PDH-ir projections forming dorsal and ventral commissures, respectively. **c)** Filled arrowheads indicate fibres that enter the central brain through the eyestalks and exit ipsilaterally via the oesophageal connectives. The scalebar in (a) represents 100 µm, and in (b-c) 200 µm. Abbreviations: a, anterior; p, posterior; LP, lateral protocerebrum; ES, oesophageal foramen. Figure panels showing the same individual are indicated with the same Es#ID.

### An extensive network of PDH-positive fibres in the central brain reveals contralateral connections between the eyestalks, as well as ipsilateral connections between the eyestalks and the periphery

While we found only a few PDH-positive cells in the central brain of krill, complex networks of PDH-positive fibres are visible throughout most parts of it (Figure 2c-e, 4a-c, and Figure 4-figure supplement 1). In the medial protocerebrum, the most prominent structures visible are PDH-positive fibre bundles, originating from the lateral protocerebrum in the eyestalks, which project through the eyestalks into the central brain where they form prominent dorsal and ventral commissures (filled and blank arrowheads in Figure 4b; Figure 4-figure supplement 1). Due to the dense arborisations in the lateral protocerebrum, the origin of these fibres could not be identified unambiguously. Parts of the commissural fibre bundles in each hemisphere of the central brain branch and project posteriorly, forming extensive arborisations in large parts of the posterior protocerebrum and tritocerebrum (Figure 4b and c). In the tritocerebrum, several PDH-positive fibres leave the central brain via the oesophageal connectives (Figure 4c). Specifically, we could trace fibres that originate in the eyestalks, extend through the eyestalks into the central brain, where they project into the ventral posterior brain and leave the central brain through the oesophageal connective of the respective hemisphere (filled arrowheads in Figure 4c), thus providing ipsilateral connections of the eyestalks and the periphery.

### PDH co-localizes with *cry2* and *per* mRNA in the eyestalks of krill

To assess the expression of the clock genes *cry2* and *per* throughout the krill eyestalks, we used RNA *in situ* hybridization on horizontal vibratome sections of the krill brain. Antibodies and RNA probes targeting additional TTFL components were tested but did not yield signals, so the analysis is restricted to the markers reported here. Due to methodological limitations, the two *in situ* hybridization probes could not be applied to the same sections; therefore, co-localization of *cry2* and *per* could not be assessed directly. However, a combination of *in situ* hybridization and antibody staining allowed indirect co-localization analysis. The *in situ* hybridization signal of both *cry2* and *per* probes shows distinct staining of two cell clusters, whose location closely resembles those of the lamina- and lobula-clusters described for PDH-positive cells above (Figure 5-figure supplement 1a-d). Negative control sections processed identically but without the addition of the RNA probes showed no detectable signal, confirming the specificity of the signal (Figure 5-figure supplement 2). Since our data show that both clock genes are expressed in the same brain regions, we next performed *cry2* and *per in situ* hybridization with subsequent PDH-immunolabeling. The double-labelling revealed that both *cry2* and *per* co-localize with all PDH-positive cells in the lamina- and lobula-cluster (filled arrowheads in Figure 5), suggesting that *cry2* and *per* are themselves co-expressed, at least in these PDH-positive neurons. Moreover, in both clusters, the co-labelling revealed additional *cry2*- and *per*-positive cells which are not PDH-positive (blank arrowheads in Figure 5), showing that the number of clock gene-expressing cells in these clusters is higher than the number of cells containing PDH.

**Figure 5:**
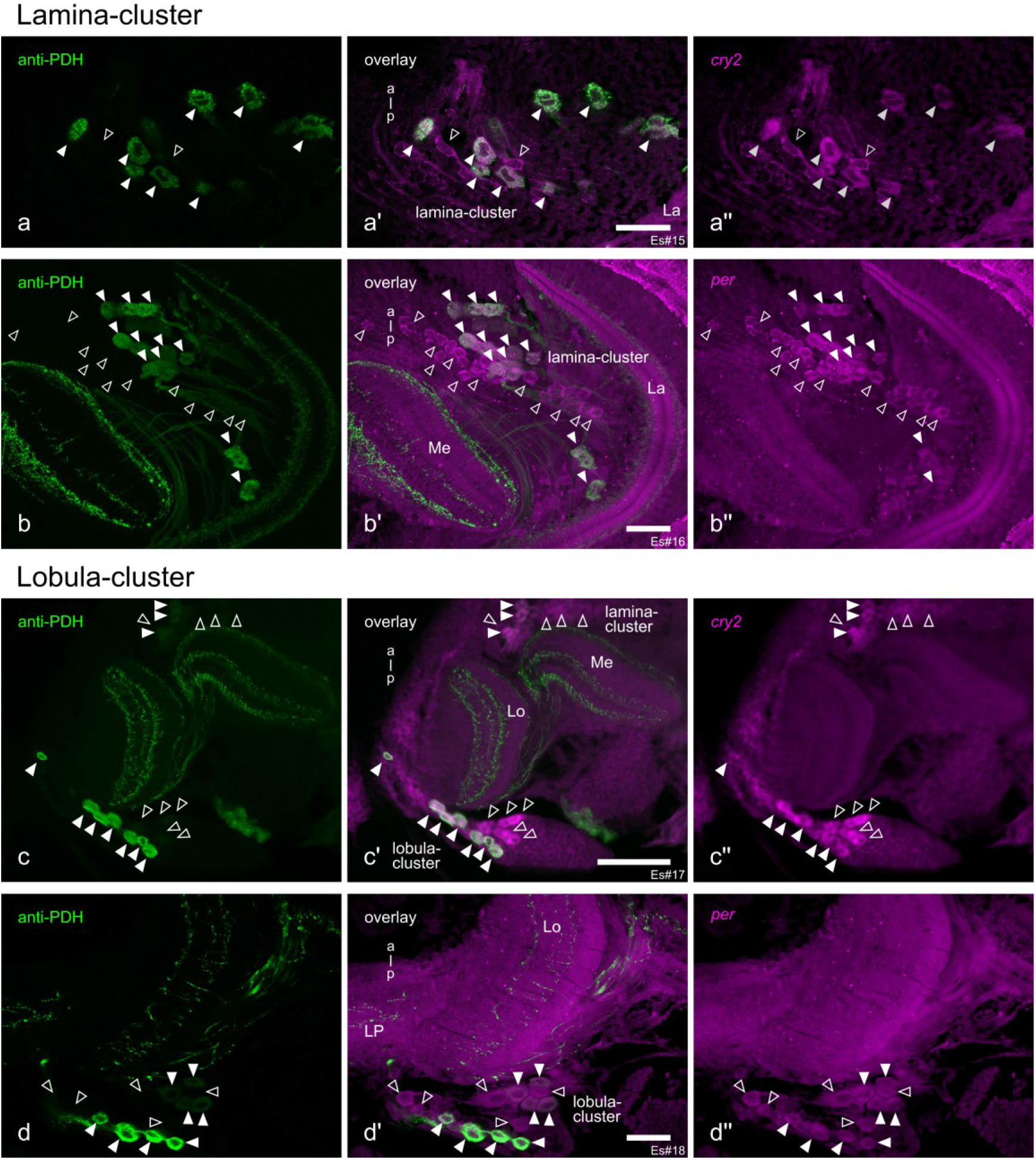
PDH-positive cells co-localize with *cry2* and *per* transcripts in the krill eyestalks. a-b’’) PDH-positive (green, a, b), as well as *cry2*- (magenta, (a’’)) and *per-*positive (magenta, (b’’)) cells in the lamina-cluster of the krill eyestalk. **a’, b’)** show co-localization of the anti-PDH and in situ hybridization signal in a subset of *cry2*- (filled arrowheads in a’) and *per-*positive (filled arrowheads in (b’)) cells. **c-d’’)** PDH-positive (green, (c, d)), as well as *cry2*- (magenta, (c’’)) and *per-*positive (magenta, (d’’)) cells in the lobula-cluster. The PDH-positive cells co-localize with a subset of the *cry2*- (filled arrowheads in (c’)) and *per*-positive (filled arrowheads in (d’)) cells, respectively. Blank arrowheads in (a-d’’) show *cry2*- and *per-*expressing cells that do not co-localize with anti-PDH. Note that methodological limitations prevented direct co-localization of *cry2* and *per* in the same sample. Abbreviations: La, lamina; Me, medulla; Lo, lobula; LP, lateral protocerebrum. Scale bars represent 100 µm. Figure panels showing the same individual are indicated with the same Es#ID.

### *cry2* and *per* are widely expressed but do not co-localize with PDH in the krill central brain

In contrast to the expression patterns in the eyestalks, *in situ* hybridization revealed extensive expression of *cry2* and *per* throughout large parts of the central brain (Figure 6a-h). The staining pattern of the two probes is largely similar. In the dorsal brain, *cry2*- and *per*-positive cells are mainly found in the posterior brain (Figure 6a and e). Further ventrally, the *in situ* hybridization signal is mostly restricted to the lateral and posterior brain, except for two clusters of cells in an anterior medial position (Figure 6b and f). Even further ventral, *cry2*- and *per*-expression is restricted to the most lateral parts of the central brain (Figure 6c and g), while in the most ventral sections, *cry2*- and *per*-positive cells are mainly found in the centre of the brain between the prominent neuropil regions of the first and second antenna, and the olfactory lobes (Figure 6d and h). Interestingly, in separate co-staining experiments, neither *cry2* nor *per* transcripts appeared to co-localized with PDH-positive cells in the central brain (Figure 6i-j’’, green arrowheads). As we focused our analysis only on the few strongly PDH-positive cells in the central brain we cannot exclude that some of the weakly PDH-ir cells might co-localize with *per* or *cry2*. However, due to the methodological limitations further staining will be necessary to unravel the expression of clock genes in these cells.

**Figure 6:**
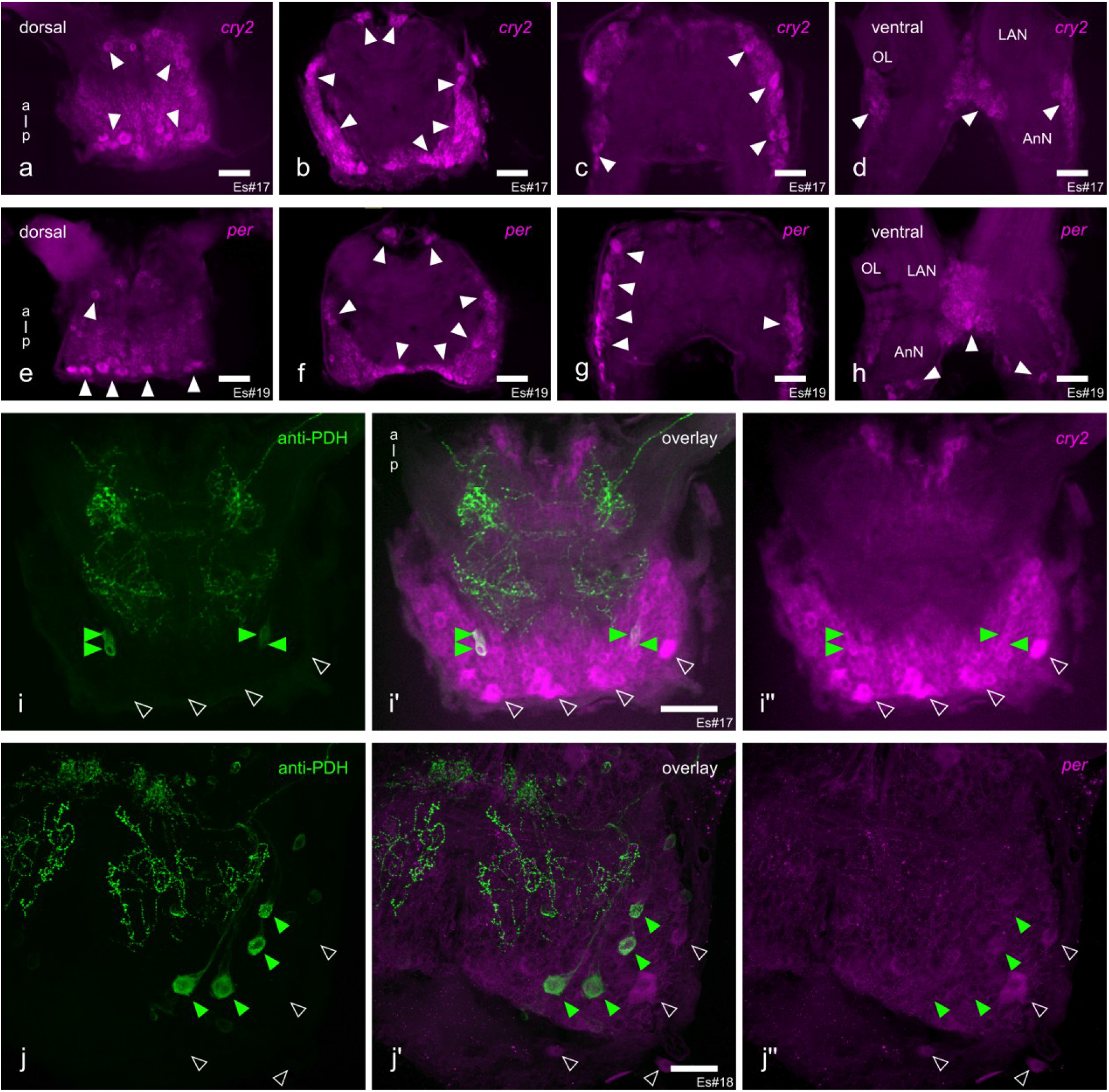
Distribution of *cry2* and *per* transcripts in relation to the PDH-network in the krill brain. a-h) Stereo microscopic images of 100 µm sections of the krill central brain showing cells expressing *cry2* (a-d) and *per* (e-h) transcripts. From left to right, images show sections progressing from dorsal to ventral regions of the central brain. Areas with high *cry2*-/ *per*-expression are marked by white arrowheads. **i-i’’)** PDH-positive neurons in the krill central brain (green arrowheads) do not co-localize with *cry2*-expressing neurons (blank arrowheads). **j-j’’)** Similarly, PDH-positive neurons (green arrowheads) do not co-localize with *per*-expressing neurons (blank arrowheads) as revealed by RNA in situ hybridization. Abbreviations: OL, olfactory lobes; LAN, lateral antenna 1 neuropil; AnN, antenna 2 neuropil. Note that methodological limitations prevented direct co-localization of *cry2* and *per* in the same sample. The scale bars represent 200 µm in (a-h) and 100 µm in (i-j’’). Figure panels showing the same individual are indicated with the same Es#ID.

## Discussion

The rhythmic adaptation of krill to its high-latitude environment is key to its success in the Southern Ocean, which in turn represents a cornerstone for the well-being of the whole krill-centred ecosystem. To better understand the mechanisms that govern daily and seasonal timing in krill, we provide a detailed description of the PDH-neuronal network and show that the PDH-positive neurons in the eyestalks also express the circadian clock genes *cry2* and *per,* suggesting that they are genuine circadian clock neurons (Figure 7). This finding represents an important advance in our aim to understand the mechanistic basis of biological timing in krill.

**Figure 7:**
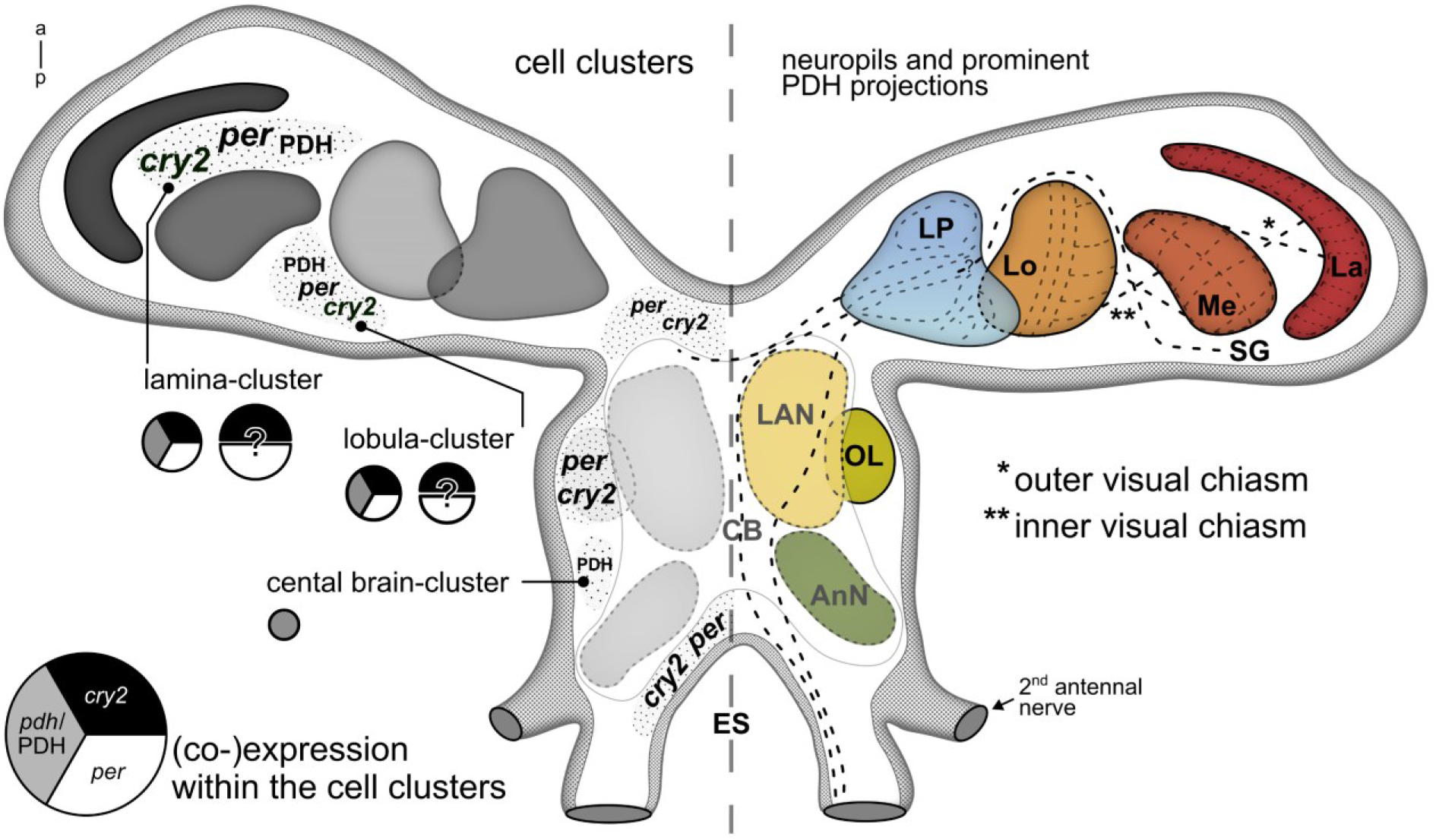
Location of putative clock neurons and neuropil regions in the brain of krill. <u>Left hemisphere:</u> Schematic representation of the general location of neurons expressing *cry2* and *per* and or *pdh*/PDH (dotted areas, label size is representing relative cell abundance). Note that weakly PDH-ir cells are excluded from this schematic. Cells expressing both, *cry2* and *per* (indirectly shown by co-expression of PDH) are considered candidate clock neurons. In the eyestalk, the location and the number of *cry2*- and *per*-expressing neurons suggest co-localization also in PDH-negative neurons, categorizing them as potential clock neurons. However, we could not investigate co-localization via *in situ* hybridization due to methodological limitations. *cry2*- and *per*-expression in the central brain was present in numerous cells, but distinct clusters could not be defined. Note that PDH-positive cells in the central brain appeared to not co-localize with *cry2*- or *per*-expressing cells. Pie charts indicate which transcripts or protein is expressed in the respective cluster (grey = *pdh*/PDH, black = *cry2*, white = *per*). The size of the pie charts indicates the relative estimated size of the respective cell cluster. Question marks indicate that co-localization of *cry2* and *per* has not been shown but is assumed based on location and numbers of cells. <u>Right hemisphere:</u> Schematic representation of the neuropils identified in the krill brain. Black dashed lines represent major PDH-positive projections. Abbreviations: La, lamina; Me, medulla; Lo, lobula; LP, lateral protocerebrum; SG, sinus gland; LAN, lateral antenna 1 neuropil; OL, olfactory lobes; AnN; antenna 2 neuropil; ES, oesophageal foramen; CB, central brain.

### The krill PDH-network shows similarities with those of other malacostracan crustaceans

#### Conserved PDH-ir clusters in the eyestalks

As nicely summarized by Meelkop & Janssen (2011), the most striking similarities found between the PDH-ir patterns in the eyestalks are clusters of PDH-ir cells between the lobula and medulla terminalis, as well as their associated fibre tracts, with the most prominent tract targeting the neurohemal sinus gland. The cell clusters between the lobula and lateral protocerebrum have been referred to as cluster C (Harzsch et al., 2009; Luna et al., 2010) or clusters 2, 4, and 5 (Chung & Webster, 2004) in previous malacostracan studies, and match the position of the cells of the lobula-cluster we described for *E. superba* in the present study. Earlier work reported that a subpopulation of neurons within this region forms T-shaped axons projecting laterally to the sinus gland and medially towards the lateral protocerebrum (Harzsch et al., 2009), a pattern consistent with our observations in krill. Although we could not unambiguously trace the full projection pattern of these neurons in krill, our data indicate that the medial branches of the lobula-cluster are likely to pass through the eyestalks and continue into the medial protocerebrum. This would match the situation described in *Homarus americanus*, where cluster C projections could be traced both laterally into the sinus gland and medially into the protocerebrum (Harzsch et al., 2009; Strauß & Dircksen, 2010).

Interestingly, studies on the ontogeny of the PDH-ir network showed that cells of cluster C are present from the earliest embryonic stages on in *Carcinus maenas* and *Homarus americanus* (Chung & Webster, 2004; Harzsch et al., 2009) and are reminiscent of the patterns in adult animals by the end of embryonic development. The early establishment of the PDH-ir network in the eyestalks has been suggested to reflect its involvement in the circadian pacemaker system, although its functional significance remains unknown (Chung & Webster, 2004; Harzsch et al., 2009).

In addition to the distinct PDH-positive clusters, we found single PDH-positive cells that project into the lateral protocerebrum. Similarly, additional single or pairs of PDH-ir cells have been identified in *Homarus americanus*, which project towards the lateral protocerebrum and continue through the eyestalk into the medial protocerebrum (termed A and B cells in Harzsch et al., 2009). In krill, however, we lost their projections in the dense arborisations of the lateral protocerebrum.

#### A distinctive PDH-ir lamina-cluster

Although the position and projection patterns of the lobula-cluster described here largely match those of cell clusters described for various crustacean species, the lamina-cluster is the largest, most prominent PDH-positive cluster we identified in krill. While it was difficult to follow the single fibres due to their frequent superposition, most of the lamina-cluster neurons appear to send projections into the lamina; only a few direct projections into the medulla have been identified. In other malacostracans, clusters of cells which primarily target the lamina have been documented (e.g., *Carcinus maenas* (Alexander et al., 2020; Mangerich et al., 1987), *Cancer productus* (Hsu et al., 2008), *Orconectes limosus* (de Kleijn et al., 1993), and *Procambarus virginalis* (Luna et al., 2010)). The lamina-associated clusters described in these species are characterised by a large number of small PDH-ir soma, distributed across the proximal side of the lamina. While similar PDH-positive cells have been found in krill, the lamina-cluster additionally contains many large PDH-positive cells distributed in a distinct, elongated cluster between the medulla and lamina. To our knowledge, a cluster with similar morphology has so far not been reported in malacostracan or other crustaceans.

### Central brain PDH-networks and commissural connections

In the central brain of krill, the most extensive PDH-ir structures appear in the medial protocerebrum but also structures in the tritocerebrum show complex immunoreactive projections, which is in agreement with other malacostracan crustaceans studied (Chung & Webster, 2004; Harzsch et al., 2009; Hsu et al., 2008; Mangerich & Keller, 1988; Sullivan et al., 2009; Wilcockson et al., 2011). In krill, several fibre bundles travel through the eyestalks, some of which form prominent commissures in the dorsal and ventral part of the anterior medial protocerebrum. Like this, they provide a direct connection between the left and right eyestalks, which is in agreement with patterns found in many malacostracan crustaceans (Chung & Webster, 2004; Harzsch et al., 2009; Hsu et al., 2008; Mangerich & Keller, 1988; Sullivan et al., 2009; Wilcockson et al., 2011) and insects (Reischig et al., 2004; Wei et al., 2010). Two to four strongly stained PDH-positive cells per hemisphere are located dorsally in the medial protocerebrum of krill and send their projections anteriorly into the medial protocerebrum, where we lost their path. The position and projection pattern of these cells resemble single PDH-ir cells found in *Carcinus maenas* and *Orconectes limosus*, which send their projections into the anterior medial protocerebral neuropil, termed optic neuropil in the original study (Mangerich & Keller, 1988), as well as in *Homarus americanus* embryos, where they have been termed group E cells, which first project anteriorly but are suggested to join axon projections from the eyestalks into the oesophageal connectives (Harzsch et al., 2009).

### Potential functional roles of PDH in krill

#### PDH-mediated pigment migration and light adaptation

Early studies found PDH, together with its counterpart red pigment concentrating hormone (RPCH), as a potent effector of pigment migration in crustaceans (Rao, 2001). In this process, PDH induces the dispersion of pigments to shield vulnerable parts of the organism against harmful radiation during the day, while RPCH reverses this process into a night-adapted state. In krill, large areas of the dorsal and lateral body are covered by erythrophores, chromatophore cells which contain red pigment granules (astaxanthin in krill), shown to rapidly expand upon light exposure or stress (Auerswald et al., 2008). Due to the rapid attenuation of light in the water column, krill are exposed to varying levels of light during their daily vertical migrations. Especially during the hours just before the morning descent, when krill are feeding typically in the upper water column, they can be exposed to elevated light levels. In addition, during midsummer, when darkness during the night is absent, but krill have a strong need to feed, they may spend extended amounts of time in illuminated waters (Cisewski et al., 2010), where protection from harmful radiation is crucial, and thus pigment dispersion during these times acts as a natural sunscreen. In turn, full contraction of the pigments will render krill almost transparent, which will reduce the risk of visual predation. Besides chromatophore dispersion, PDH was found to induce movement of distal pigment cells in the eye, which puts it into a light-adapted state (Fernlund, 1976; Rao & Riehm, 1993; Verde et al., 2007). In addition, in crayfish, PDH impacts the electrical response of photoreceptor cells to light, and this response depends on the circadian phase (Solis-Chagoyan et al., 2012; Verde et al., 2007). Both processes can modulate the retinal light signal and are likely under circadian control. While light-adaptation mechanisms, such as those mentioned above, have not been studied systematically in *E. superba*, circadian rhythms of visual sensitivity measured by electroretinogram have been reported for the closely related arctic krill *Thysanoessa inermis* (Cohen et al., 2021), suggesting that a similar regulation could be present in *E. superba*, too.

#### PDH as a candidate circadian output factor

Although originally characterized as chromatophore-regulating hormones, both PDH and RPCH have since been found to occur and act beyond classical neurohemal systems and are now considered multifunctional neuropeptides with additional neuromodulatory roles. RPCH has been linked to the modulation of reproductive processes (Fingerman, 1997; Jayasankar et al., 2020; Sarojini et al., 1995), locomotor activity (Sherff & Mulloney, 1991), and the digestive system (Nusbaum & Marder, 1988), suggesting that it functions not only as a circulating hormone but also as a local signalling molecule within the Crustacean nervous system.

In contrast, PDF, the insect homologue of PDH, is best known as an output factor of the insect circadian clock, where it functions as a neurotransmitter or neuromodulator and co-localizes with a subset of clock neurons (Helfrich-Förster, 1995). Although the functional role of PDH as an output factor of the circadian clock has yet to be demonstrated in crustaceans, its direct homology to insect PDF and the general similarities in neuronal localization and projection patterns strongly suggest this. Furthermore, its involvement in regulating rhythmic physiological processes supports this interpretation. Interestingly, in contrast to insects, crustaceans have been shown to express several β-PDH isoforms, which are thought to differ in their function as neurohormone and/or neurotransmitter and thus in the regulation of the various processes described above (Alexander et al., 2020; Hsu et al., 2008; Klein et al., 1994; Mayer et al., 2015; Rao, 2001). The β-PDH antibody applied here has been shown to reliably detect two isoforms (PDH-1 and PDH-3), which are thought to cover both neurohormonal and neuromodulatory roles in other crustaceans (Alexander et al., 2020; Hsu et al., 2008). Interestingly, the PDH antibody was shown to exhibit differential reactivity to the various isoforms investigated (Alexander et al., 2020), which could explain the variations in staining intensity observed in some of the PDH-ir neurons in krill. Although the presence of PDH isoforms and their specific function as neuromodulators or -hormones has not yet been studied in krill, this represents an interesting subject for future research into the krill clock and its outputs.

### Candidate clock neurons are located in the eyestalks of krill

Due to its conserved sequence and the availability of potent antibodies, the location of PDH-ir neurons has been described in several crustaceans. But insights into the distribution of clock gene-and protein-expressing cells and their co-localization with putative output factors like PDH are scarce, and detailed descriptions, comparable to those of the insect clock system, are missing. However, to characterize the neuronal architecture of the circadian clock, the localization of its molecular components and their co-localization with output factors are prerequisites.

In crustaceans, most information is available from studies on crayfish using antibodies against *Drosophila melanogaster* clock proteins. In *Procambarus clarkii*, immunolabeling identified CRY-positive cells in the eyestalks and the anterior medial protocerebrum (Fanjul-Moles et al., 2004). Additional work using antibodies against PER, TIM, and CLK revealed their widespread expression in anterior and lateral central brain cell clusters, as well as in the eyestalk (Escamilla-Chimal et al., 2010). In agreement with that, *Cherax destructor* expresses dCRY in cells of the same anterior central brain cluster, known as cluster 6 in crustaceans (Sullivan et al., 2009). Immunocytochemical analyses in *Procambarus clarkii* detected PER in retinal photoreceptors and in cell bodies of the lamina. However, other regions were not investigated in this study (Arechiga & Rodriguez-Sosa, 1998). In *Homarus americanus*, PER cycling has been reported at the tissue level in the eyestalks, with no signal detected in central brain extracts (Grabek & Chabot, 2012), while immunolabeling in the crab *Cancer productus* revealed CYC/PDH co-localization in a cluster 6 cell in the protocerebrum (Beckwith et al., 2011).

Outside of the Decapoda, clock genes or proteins have been localized in the marine isopods *Eurydice pulchra* and *Parhyale hawaiensis* (Oliphant et al., 2025; Zhang et al., 2013), and the terrestrial isopod *Armadillidium vulgare* (Fouda et al., 2010). In *Eurydice pulchra* and *Parhyale hawaiensis*, transcripts of the clock genes *cry2*, *per,* and *tim* are restricted to the protocerebrum, where distinct clusters of clock neurons appear to be responsible for the control of circatidal and circadian rhythmicity (Oliphant et al., 2025). In *Armadillidium vulgare*, few neurons expressing CLK, CYC, and PDF were localized in the eyestalks between the medulla and lobula, as well as laterally in the suboesophageal ganglion (Fouda et al., 2010).

In summary, in crustaceans, clock components are most commonly found in both the eyestalks and the central brain, associated with the visual neuropils and the protocerebrum, respectively. This organization is consistent with the general architecture of the insect circadian system (reviewed in Helfrich-Förster et al., 1998; Numata et al., 2015) and might represent the organizational principle of the clock network in arthropods in general.

Using *in situ* hybridization together with immunohistochemistry, we found two distinct cell clusters in the eyestalk. While the number of *cry2-/per-*expressing cells was higher than the *pdh*-expressing cells, all PDH-positive cells in the eyestalk clusters showed co-localization with either *cry2* or *per*, strongly suggesting a functional role of PDH in the krill circadian clock. In contrast to the expression in the eyestalks, extensive expression of *cry2* and *per* transcripts was detected throughout large parts of the krill central brain and did not co-localize with anti-PDH. Detailed studies from several insect species show that clock neurons in the central brain are typically organized into specific clusters, most commonly found in the dorsal protocerebrum (reviewed in Helfrich-Förster et al., 1998; Numata et al., 2015). The widespread expression of *cry2* and *per* throughout most parts of the central brain of krill is in stark contrast to the general pattern found in insects, as well as its distinct expression pattern found in the krill eyestalks. Because of methodological limitations, we could not verify if *cry2* and *per* are co-expressed in these cells or if the transcripts are present in distinct cells. The broad expression might indicate that the clock transcripts in the central brain of krill may fulfil other or additional purposes unrelated to the circadian clock. Indeed, non-clock-related functions of clock genes have also been reported in insects (Beaver et al., 2003; Tobback et al., 2011; Tobback et al., 2012). Nevertheless, we cannot currently rule out the possibility that the *cry2*- and *per*-positive neurons in the central brain of krill are additional true clock neurons.

In summary, our results support the conclusion that the clock gene-expressing neurons located in the eyestalks represent a pacemaker of the circadian clock in krill. However, due to the current lack of antibodies against krill clock proteins, our analysis of core clock components is limited to the transcript-level, which might not fully represent the protein level. In addition, methodological limitations currently prevent us from investigating daily and circadian changes in clock gene transcript levels. In an earlier study on krill, Mazzotta et al. (2010) reported subtle daily changes in CRY2 levels by Western blot on extracts of whole krill heads, suggesting that CRY2 might cycle at the protein level in krill. Future studies may build on our findings to localize clock proteins and quantify their temporal changes at the cellular level.

### Potential homology to the insect circadian clock network

Across the tree of life, considerable variability is observed in the presence and role of single clock genes and proteins (Kotwica-Rolinska, Chodakova, et al., 2022; Tomioka & Matsumoto, 2010), the use of retinal or extraretinal entrainment pathways (Helfrich-Förster, 2019; Yoshii et al., 2015), and the exact location and projections of clock neurons (Helfrich-Förster, 2004, 2017; Helfrich-Förster et al., 1998; Numata et al., 2015). However, its general molecular structure and gross neuronal architecture are conserved at least across insects (Helfrich-Förster et al., 1998; Numata et al., 2015). In terms of the neuronal architecture, a unifying feature in the distribution of the circadian clock appears to be the separation into dorsal (DN) and lateral (LN) clock neurons. While the DN are found in the dorsal central brain, typically anterior in the medial protocerebrum, the LN are closely associated with neuropils of the light signalling pathway (Helfrich-Förster et al., 1998; Numata et al., 2015).

Due to the well-known function of PDF as a clock output factor in insects (reviewed in Hermann-Luibl & Helfrich-Förster, 2015; Shafer & Yao, 2014) and based on the location and projection patterns of PDH-ir neurons, several studies have discussed potential homologies of crustacean PDH-ir clusters to the well-studied insect circadian clock architecture (reviewed in Strauß & Dircksen, 2010). In the following, we will briefly outline similarities between clock neurons and PDH-ir structures found in krill and insects.

Although absent in flies, the most lateral PDH-ir clusters are those associated with the lamina, as found in locusts, crickets, and cockroaches (Homberg & Prakash, 1996; Homberg et al., 1991; Nässel et al., 1991; Wei et al., 2010). In these insects, the neurons are organized in distinct ventral and dorsal clusters at the proximal side of the lamina. Cockroaches lack the light-sensitive dCRY, and entrainment of the clock depends on the compound eyes. The PDF-positive lamina neurons are thought to be involved in this pathway (reviewed in Helfrich-Förster, 2019; Stengl et al., 2015). In krill, we identified a PDH-positive cluster of candidate clock neurons with large somata whose projections mainly target the lamina. In addition, we observed extensive PDH-positive connections between the lamina and medulla, as well as the medulla and lobula, which could at least partly originate from cells of the lamina-cluster. In addition, we showed that the PDH-positive cells of the lamina-cluster co-localize with transcripts of *cry2* and *per*, which are thought to function as transcriptional repressors in the krill core clock (Biscontin et al., 2017). Thus, the clock neurons of the lamina-cluster could function as an input pathway of the retinal light signal to the circadian clock. At the same time, they could provide a mechanism to modulate the incoming light signal in a circadian fashion. Additionally, they could also contain circadian clock neurons that control behavioural rhythms, as suggested for the cricket, *Gryllus bimaculatus* (Abe et al., 1997; Tomioka & Chiba, 1986, 1992) and the New Zealand weta *Hemideina thoracica* (Waddell et al., 1990), in which the main circadian pacemaker neurons were traced to neurons located between the lamina and medulla. Although up to now we can only speculate on the functional role of the lamina clock neurons in krill, the general size of their compound eyes, the localization of putative pacemakers in the eyestalks and their overall light-centred lifestyle collectively indicate a high priority towards light processing and signalling, which could explain the prominent lamina-cluster in krill.

Another cluster of PDH-positive clock neurons is located between the medulla and lobula in krill, which closely resembles cluster C neurons in other crustaceans (reviewed in Meelkop et al., 2011).

Based on their characteristic position between medulla and lobula, as well as their arborization patterns, projecting into the medial protocerebrum and the contralateral eyestalk they have been thought to be homologues of LN found closely associated with the accessory medulla in several insects, including flies, bees, cockroaches, and aphids (Beer et al., 2018; Colizzi et al., 2023; Helfrich-Förster & Homberg, 1993; Wei et al., 2010). The accessory medulla is a key centre of the circadian clock, which integrates photic input and the signals from various clock neurons to control rhythmic output (reviewed in Helfrich-Förster, 2019; Stengl et al., 2015).

As shown for several decapod crustaceans, a subset of PDH-positive neurons of the lobula-cluster seems to terminate in the sinus gland, and in krill, these cells express *cry2* and *per*, strongly suggesting that they form a part of the clock network. The sinus gland is a major neurohemal centre in crustaceans, serving as the primary release site for a diverse set of regulatory hormones (reviewed in Simoes et al., 2024). Neuropeptides such as PDH, RPCH, molt-inhibiting hormone, gonad-inhibiting hormone, and members of the crustacean hyperglycemic hormone family are secreted from the sinus gland into the haemolymph, where they regulate processes such as light sensitivity and chromatophore-mediated colour change, moulting, vitellogenesis, maturation, and osmoregulation. Further, rhythmic hormone release from the sinus gland is thought to drive circadian rhythms in the response to light, which depends on PDH (Hernández-Falcón et al., 1987; Moreno-Sáenz et al., 1987; Moreno-Sáenz et al., 2005). In addition, daily rhythms in PDH-immunoreactivity of the SG were found in *Cancer productus* (de la Iglesia & Hsu, 2010). Like many other zooplankton species, krill regularly performs DVM in the water column, which represents an adaptation to the daily change in predation pressure, driven by the daily light-dark cycle (Bahlburg et al., 2023; Gastauer et al., 2021; Hobbs et al., 2021). Supporting a role for endogenous timing mechanisms in this behaviour, we recently demonstrated that changes in krill swimming activity are under circadian clock control (Hüppe et al., 2025). Although not explicitly studied in *E. superba*, adaptations of the visual system in response to the day-night cycle have been shown for other daily migrators (Cohen et al., 2021; Deleo & Bracken-Grissom, 2020). Besides daily rhythms like DVM, krill show pronounced physiological adaptations in response to the strong seasonality of the Southern Ocean environment. This includes active reproduction, development, and growth in summer, and metabolic reduction and lipid accumulation, as well as halted development in winter (Kawaguchi et al., 2007; Meyer et al., 2010). Most interestingly, the sexual maturity cycle, closely linked to their seasonal reproduction, is under photoperiodic control (Höring et al., 2018). Recent studies could show that the circadian clock provides daylength information to time seasonal events and that PDF plays a key role in signalling photoperiodic information to downstream neuroendocrine centres (reviewed in Helfrich-Förster, 2024; Hidalgo et al., 2023; Nagy et al., 2019). In the context of clock-dependent timing of daily and seasonal physiology and behaviour, the direct connection between PDH-positive putative clock neurons and the sinus gland in krill thus provides a promising target for future studies on the neuronal basis of daily and seasonal adaptation.

Temporal synchronization at both daily and seasonal timescales is central to the success of krill in the Southern Ocean. Over the past decade, growing evidence has highlighted the role of endogenous timing mechanisms in shaping this temporal adaptation (Höring et al., 2018; Hüppe et al., 2025; Piccolin, Meyer, et al., 2018; Piccolin et al., 2020; Piccolin, Suberg, et al., 2018). At the same time, reliance on internal clocks may constrain responses under conditions of rapid environmental change (Helm et al., 2013; Visser & Gienapp, 2019). However, as in most crustaceans, chronobiological research in krill has long remained predominantly phenomenological, providing valuable descriptive insights but leaving many mechanistic questions unresolved. In this study, we present findings that offer a foundational framework for dissecting the underlying mechanisms that drive rhythmic processes in krill. By establishing this mechanistic basis, our work opens the door to a deeper understanding of how these rhythms shape physiological functions, behaviour, and ultimately the adaptive strategies of krill. Together, these advances represent an important step towards a more comprehensive and mechanistic understanding of krill biology and ecology.

## Material and Methods

### Animal collection and fixation

Adult krill (*E. superba*) of mixed sexes were sampled from the Atlantic sector of the Southern Ocean, north of the South Orkney Islands (60.1°S, 45°W) during austral late summer and autumn (8^th^ February to 21^st^ March 2022). Krill were sampled from the continuous fishing system of the krill fishing vessel Antarctic Endurance at regular intervals over the daily cycle. Krill were pumped aboard via a vacuum hose attached to the cod end of a commercial trawl, while the vessel trawled at a speed of 1.5 to 2 knots. On deck, the krill were separated from the water on a metal grate, and specimens were sampled from there. The collected krill were kept in surface seawater at 1 °C in constant darkness and were further processed within <20 min. After sampling, sex and length were determined under a stereo microscope before decapitating and fixing their heads in 4% paraformaldehyde (PFA) stabilized with Hexamethylenetetramine. Additionally, whole krill individuals were fixed for whole central nervous system preparations. Samples were stored at 4°C in PFA until further processing, which took place between June 2023 and October 2025. Importantly, we did not observe changes in staining quality over time, even if storage time in PFA varied throughout the project.

### Sample dissection and preparation

PFA-fixed krill heads were washed three times while whole krill were washed five times in phosphate-buffered saline (PBS) at room temperature (RT) for ten minutes before dissection in PBS containing 0.5% Triton X-100 (PBST 0.5%). During dissection, the cuticle and muscle tissue were removed to reveal the nervous tissue of the central brain and eyestalks. To facilitate subsequent vibratome sectioning, the retina and ommatidia, especially the crystalline cones of the compound eyes, were removed.

For *in situ* hybridization, either whole brains or vibratome sections were dehydrated in a methanol series (30%, 50%, 70%, 90%, and 100% methanol in PBS) for about 20 minutes each. After an overnight incubation at −20°C in 100% methanol, the samples were rehydrated by reversing the methanol series. After rehydration, the samples were kept in PBS at 4°C.

For antibody staining, antigen retrieval was carried out by incubating the brains in citrate buffer (0.01 M citric acid monohydrate, pH = 6.0 with NaOH) for 30 minutes at 80°C. Afterward, the brains were washed three times in PBST 0.5% for 10 minutes and stored at 4°C.

Finally, for both antibody staining and RNA *in situ* hybridization, the brains were embedded in 7% agarose (Sigma, A9539) and sectioned into 100-200 µm slices in ice-cold PBS, using a vibratome (Leica VT1000S; 0.225 mm/s at 50 Hz).

### Immunohistochemistry

Horizontal vibratome sections of krill brains were incubated in a blocking solution (5% normal goat/donkey serum, NGS/NDS in PBST 0.5% + 0.02% NaN_3_) for at least three hours at room temperature (RT) or overnight at 4°C. After blocking, the samples were incubated at 4°C for two nights in a primary antibody solution containing 5% NGS/NDS in PBST 0.5%, 0.02% NaN_3_, and the primary antibodies in the specified concentration. For labelling PDH-positive neurons, a polyclonal antibody raised in rabbit against synthetic *Uca pugilator* β-PDH (Dircksen et al., 1987; Wilcockson et al., 2011) was used (1:4000). Anti-β-PDH was kindly provided by David Wilcockson (Aberystwyth University, Wales). For neuropil labelling, a monoclonal anti-Synapsin (1:50) antibody (DSHB Cat# 3C11, RRID: AB_528479) was used (Klagges et al., 1996). The incubation with the primary antibody solution was followed by six washes with PBST 0.5% at RT for 20 minutes each. Then, the sections were incubated overnight in a secondary antibody solution containing 5% NGS/NDS in PBST 0.5%, 0.02% NaN_3_, and the corresponding secondary antibodies at a concentration of 5 µg/mL. Highly cross-adsorbed secondary antibodies raised in either goat or donkey, and conjugated with Alexa Fluor 488, 555, or 647, were purchased from Thermo Scientific. As a reference staining, the nuclei were counterstained with Hoechst 33342 (10 µM) during the secondary antibody incubation. Vibratome sections were washed five times with PBST 0.5% for 10 minutes before embedding. The sections were arranged on an object slide and mounted in Vectashield antifade mounting medium (Vector Laboratories; H-100) under coverslips sealed with FixoGum (Marabu GmbH). The samples were stored in the dark at 4°C until imaging. For each staining at least five replicates were used.

### *In situ* hybridization

#### Probe generation

To map the expression of clock gene transcripts in krill, RNA probes targeting *E. superba cry2* and *per* mRNA were synthesized from krill cDNA.

Krill cDNA was obtained from whole head RNA extracts using krill collected from the Southern Ocean, as described above. After sampling, whole krill heads were fixed in RNA*later*™ stabilizing solution (Thermo Scientific) for 24 hours at 4°C and stored at −80°C until further processing. The heads were homogenized in TRIzol™ Reagent (Thermo Scientific), and total RNA was extracted using a combination of chloroform-based phase separation followed by purification with the Direct-zol Miniprep Plus Kit (Zymo Research). The RNA quality and quantity were assessed photometrically using a NanoDrop (Thermo Scientific) and via on-chip gel electrophoresis using a 2100 Bioanalyzer (Agilent Technologies). Subsequently, cDNA was synthesized using the RevertAid H Minus First Strand cDNA Synthesis Kit (Thermo Scientific).

Krill cDNA (1.3 ng/µL) was used with the Phusion™ High-Fidelity PCR Kit (New England Biolabs) and the corresponding primers for each gene (0.5 µM), as detailed in Supplementary Table 1, following the manufacturer’s protocol. The PCR products were validated on a 1% agarose gel using gel electrophoresis and purified using the MSB-Spin PCRapace & Fragmet Clean Up Kit (Stratec).

Next, PCR products were cloned to obtain plasmid templates. 15 ng of blunt-end amplicons were ligated into the vector (pJET1.2/blunt) using the CloneJET PCR Cloning Kit (Thermo Scientific). Vectors with inserts were then transformed into *E. coli* using Stellar™ Competent Cells. 5 µL of each ligation mixture were added to the competent cells (50 µL) and incubated on ice for 20–30 minutes. Then, the cells underwent a one-minute heat shock at 42 °C, with brief mixing halfway through. After adding 200 µL of Luria–Bertani (LB) medium, the cells were allowed to recover for one hour at 37 °C. For each construct, the transformed cultures were plated on LB-ampicillin agar and incubated for 16–18 hours at 37 °C. After incubation, single colonies were picked and incubated with 4 mL of LB medium containing ampicillin at 37 °C for 16 hours on a shaker.

To extract the plasmid DNA from each culture, 4 mL of the culture was pelleted by centrifugation and used with the Plasmid Miniprep Kit (New England Biolabs), following the manufacturer’s protocol. The subsequent plasmid DNA was validated by restriction digest (37 °C, 1 h) with XbaI and XhoI (FastDigest restriction enzymes, Thermo Scientific), followed by 1 % agarose gel electrophoresis. Plasmids were quantified using a Nanodrop and verified by Sanger sequencing (Mix2Seq, Eurofins). Only constructs containing the correct insert in antisense orientation were used for probe synthesis.

The plasmids (1-2 µg) verified by sequencing were linearized with XbaI at 37 °C, 1 h, checked on a 1 % agarose gel by gel electrophoresis, and then purified using the MSB-Spin PCRapace & Fragment Clean-Up kit.

Digoxigenin (DIG)-labelled RNA probes were synthesized from 700–1000 ng of linearized plasmid using T7 RNA polymerase and a DIG RNA labelling mix (Roche) in a 20 µL reaction and incubated for two hours at 37 °C, according to the manufacturer’s protocol. Reactions were terminated with 2 µL of ethylenediaminetetraacetic acid (EDTA, 0.2 µM, pH 8). The transcripts were then purified using LiCl/ethanol precipitation (2.5 µL, 4 M LiCl), followed by a 70% ethanol wash. The samples were dried and resuspended in 50 µL nuclease-free water. To each resuspended probe, 200 µL of hybridization buffer (Hyb) was added to dilute the probes for storage at −80 °C. Hyb contained 50% formamide, 0.1 µg/mL tRNA and 0.1 µg/mL ssDNA, 2% blocking solution for nucleic acid hybridization (Roche), and 0.05 µg/mL heparin in 5x saline-sodium citrate buffer pH 7 and 0.1% Triton-X 100.

#### RNA in situ hybridization procedure

Hybridized RNA probes were labelled with an anti-DIG antibody and visualized using enzymatic reactions with alkaline phosphatase (AP) with either BCIP/NBT (Thermo Scientific) or Fast Red (Sigma-Aldrich). In addition to the coloured precipitate produced by the AP reaction, Fast Red can be detected by excitation at 570 nm.

The tissue permeability and RNA accessibility were increased by incubating the vibratome sections for 10 minutes in PBST 0.3%, followed by a 10-minute treatment with proteinase K (10 µg/mL). Proteinase K activity was stopped by adding glycine (40 µg/mL) and subsequent incubation at RT for 10 minutes. The samples were refixed with 4% PFA in PBST 0.3% for one hour at RT, followed by three PBST 0.3% washes for 10 minutes each. Unless noted otherwise, all following incubation steps and reagents used during *in situ* hybridization were carried out and applied at 48 °C in a water bath. The samples were incubated for 10 minutes in a solution of 50% Hyb in PBST 0.3%. Next, the samples were prehybridized in Hyb for one hour. *Cry2* and *per* RNA probes were diluted to their final concentration with Hyb and allowed to hybridize with the samples’ mRNA for at least 16 hours. Afterward, the sections were washed for 20 minutes in Hyb followed by 50% Hyb in PBST 0.3%. Finally, the samples were washed with PBST 0.3% five times for five minutes each. The RNA *in situ* hybridization was followed by antibody staining to detect the DIG-labelled RNA probes. The blocking and primary antibody incubation were performed as described for immunohistochemistry. For samples subjected to RNA *in situ* hybridization before antibody staining, the primary antibody solution additionally contained AP-conjugated anti-DIG antibody raised in sheep (1:5000; Fab-fragments; Roche). Before the incubation with the secondary antibody solution, the AP reaction was initiated. The samples were washed three times for 10 min and two times for 20 min in PBST 0.5% and then five times for 10 min with a phosphate-free AP buffer at room temperature. AP buffer contained 1.2% TRIS, 0.6% NaCl, and 0.1% MgCl_2_ in water (w/v); the pH was adjusted to 9.5 with HCl. For the AP reaction, the samples were incubated in the dark in a staining solution either for BCIP/NBT or Fast Red and monitored closely until tissue coloration was observed (10 minutes to four hours). After successful detection of the RNA probes, the reaction was stopped by at least four 10-minute washes with PBST 0.5%. Afterward, the antibody staining went on with a second blocking step followed by the incubation with the secondary antibody solution. To rule out that the detected signal originates from endogenous AP activity or binding of the anti-DIG antibodies to epitopes other than the labelled RNA probes, negative controls were conducted in which the RNA probes were substituted by the same volume of Hyb. Else the procedure was kept the same. For each staining at least five replicates were used.

### Image acquisition

Images were acquired with a Leica TCS SP8 confocal microscope (Leica Microsystems, Wetzlar, Germany) equipped with a hybrid detector. A white light laser (Leica Microsystems, Wetzlar, Germany) was used for excitation. We used a 20-fold glycerol immersion objective (0.73 NA, HC PL APO, Leica Microsystems, Wetzlar, Germany) for scans of the vibratome sections. We obtained 8-bit or 16-bit confocal stacks with a maximal voxel size of 1.14 x 1.14x 2.0 μm to 0.76 x 0.76 x 2 μm for overview recordings and 0.76 x 0.76 x 2 μm to 0.23 x 0.23 x 2 μm for details. The optical section thickness was always 3.12 μm. Image tiles for overview images were recorded with 1024 x 1024 pixels and 2048 x 2048 pixels for detail images using LasX (3.5.7.23225) with the Navigator Plugin. Image tiles were recorded with 10% overlap, and overview images of the whole vibratome section were stitched afterward in LasX with Mosaic Merge setting “blend” to “statistic”. If the signal loss was too strong along the z-axis, a z-compensation was used by increasing the laser power with increasing imaging depth.

For widefield microscopy, a Leica M165FC fluorescent microscope (Leica Microsystems, Wetzlar, Germany) and a Nikon SMZ1270i (Nikon, Minato, Japan) were used. Images were taken with LAS AF and NIS-Elements software, respectively. If necessary, images were stitched using the Pairwise Stitching Plugin (Preibisch et al., 2009) for Fiji (Version 2.16.0; Schindelin et al., 2012). For brightfield and fluorescent scans of the total nervous system of krill, the preparation was embedded in 2% Low Melt Agarose in distilled water, using a customized silicon spacer (thickness: 1.5 mm), which was fixed with glycerol gelatine to a microscope slide. The nervous system was imaged using a Leica Inverted Microscope Thunder Imager (Leica Microsystems, Wetzlar, Germany). The microscope was equipped with Thunder deconvolution. The samples were imaged as Z-stacks (minimum voxel size: 1.3047 x 1.3047 x 62.6722 µm). Images were only adjusted linearly for brightness and contrast, using Fiji.

### 3D Reconstruction

For the reconstruction of the brain, five vibratome sections of the same brain were scanned as a tile scan. Each vibratome section was merged in LasX. The merged sections were opened in Fiji, and any remaining signal loss along the z-axis was corrected for using the Stack Contrast Adjustment Plugin (Capek et al., 2006). The adjusted images were opened with the Big Stitcher Plugin (Hörl et al., 2019). The vibratome sections were manually aligned using nuclear and neuropil staining as a reference. The aligned sections were exported as a single stack and the neuropils were reconstructed using TrakEM2 (Cardona et al., 2012). Afterward, the reconstructed neuropils were exported as labels, and Fiji 3D Viewer was used to generate 3D objects. The 3D objects were then imported into Blender (5.0; Blender Development Team, 2025), and the surfaces were smoothed to remove artefacts. For the 3D visualization of the PDH-network, the Microscopy Node Plugin for Blender (2.2.7; Gros et al., 2025) was used. For the 3D rendering, the PDH-fibre network was extracted from the background using a custom macro (Reinhard et al., 2025).

### Cell Counting

To determine the number of PDH-positive cells per cluster three whole brain datasets of mixed sexes (Es#5, Es#11, Es#20; see supplementary table 2) were reconstructed from horizontal vibratome sections as described above for the 3D reconstructions. Each cell was marked in the section its nucleus diameter was the largest using the polygon selection tool in Fiji. Based on staining intensity (compared to cells within the same focal plane), cell body size and their anatomical position in the brain the cells were assigned to the different cell clusters. Only cells with clear anti-PDH staining were considered. Cells that showed weak staining were labelled for reference but not further considered in the analysis as described in the results section. Using the “measure” function in Fiji the maximal ferret diameter as an estimate of the cell size and the cell coordinates were obtained. To obtain a measure of cluster size the number of cells per cluster and hemisphere was determined for each brain. Afterwards, the average, maximal and minimal cluster size was calculated across all hemispheres. As a measure of cell size and cell size variability, the average diameter and its standard deviation as well as the maximal and minimal diameter per cluster and hemisphere was determined for each brain. Afterwards, the average cell diameter and average maximal and minimal diameter were calculated per cluster across all hemispheres. As measure of cell size variability, the average standard deviation of the mean cell diameter was used for each cluster.

### Limitations of this approach

The lamina- and lobula-cluster extended over several sections. Hence, a reliable cell count was only possible in whole-brain-datasets. However, whole-brain-datasets at sufficiently high resolution were only feasible for a small number of samples due to the long scanning durations and high data volume. Additionally, cells might have been lost or damaged while sectioning or during dissections. Accordingly, the cell numbers remain a rough estimate of the cluster size.

### Nomenclature

In this manuscript we follow the neuroanatomical nomenclature laid out by Sandeman et al. (1992) and Richter et al. (2010), with modifications adopted from Loesel et al. (2013), Schmidt (2016), Krieger et al. (2020), and Harzsch & Krieger (2021). Specifically, we use the terms “lamina”, “medulla”, and “lobula” instead of the older terms “lamina ganglionaris”, “medulla externa”, and “medulla interna”, and collectively refer to them as “visual neuropils”. Further, we use the terms “inner” (iCh) and “outer visual chiasm” (oCh) as suggested by Krieger et al. (2020). We refer to the brain region comprising the medulla terminalis and the hemiellipsoid body, the equivalent of the insect mushroom body (Harzsch & Krieger, 2021; Strausfeld, 2020), collectively as “lateral protocerebrum” as we lack a marker to reliably distinguish between them. The brain excluding the visual neuropils and lateral protocerebrum is referred to as “central brain”.

## Acknowledgement

We thank the captain and crew of the Antarctic Endurance for their support during the field campaigns, and Aker BioMarine for their logistical assistance. We are grateful to Ryan Driscoll and Dominik Bahlburg for their help with krill sampling, and to Giulia Manoli, Moritz Bestman, and Gabriel Möller for their support during krill dissections and image acquisition. We would like to thank David Wilcockson for generously providing the PDH antibody. C.H.F. and L.H. were supported by the German Research Foundation in the framework of the priority program ‘Antarctic research with comparative investigations in Arctic ice areas’ SPP 1158 (DFG; FO 207/17-1 to C.H.F.). L.H. was further supported by the Helmholtz Association within the framework of the Initiative and Networking Fund under the Helmholtz Excellence Network Phase 2 project EXNET-02-26 and through funding by the Deutsche Forschungsgemeinschaft (DFG, German Research Foundation) under Germany’s Excellence Strategy - EXC 3051/1 „NaviSense“ – project number 533653176“. We also acknowledge funding from the DFG for the Leica TCS SP8 microscope (251610680, INST 93/809-1 FUGG) as well as the Leica THUNDER Imager (512024179, INST 93/1114-1 FUGG). We would further like to thank the Imaging Core Facility of the Biocentre for the use of the Nikon SMZ1270i.

## Supplementary Material

## Figure Supplements

**Figure 2 figure supplement 1:**
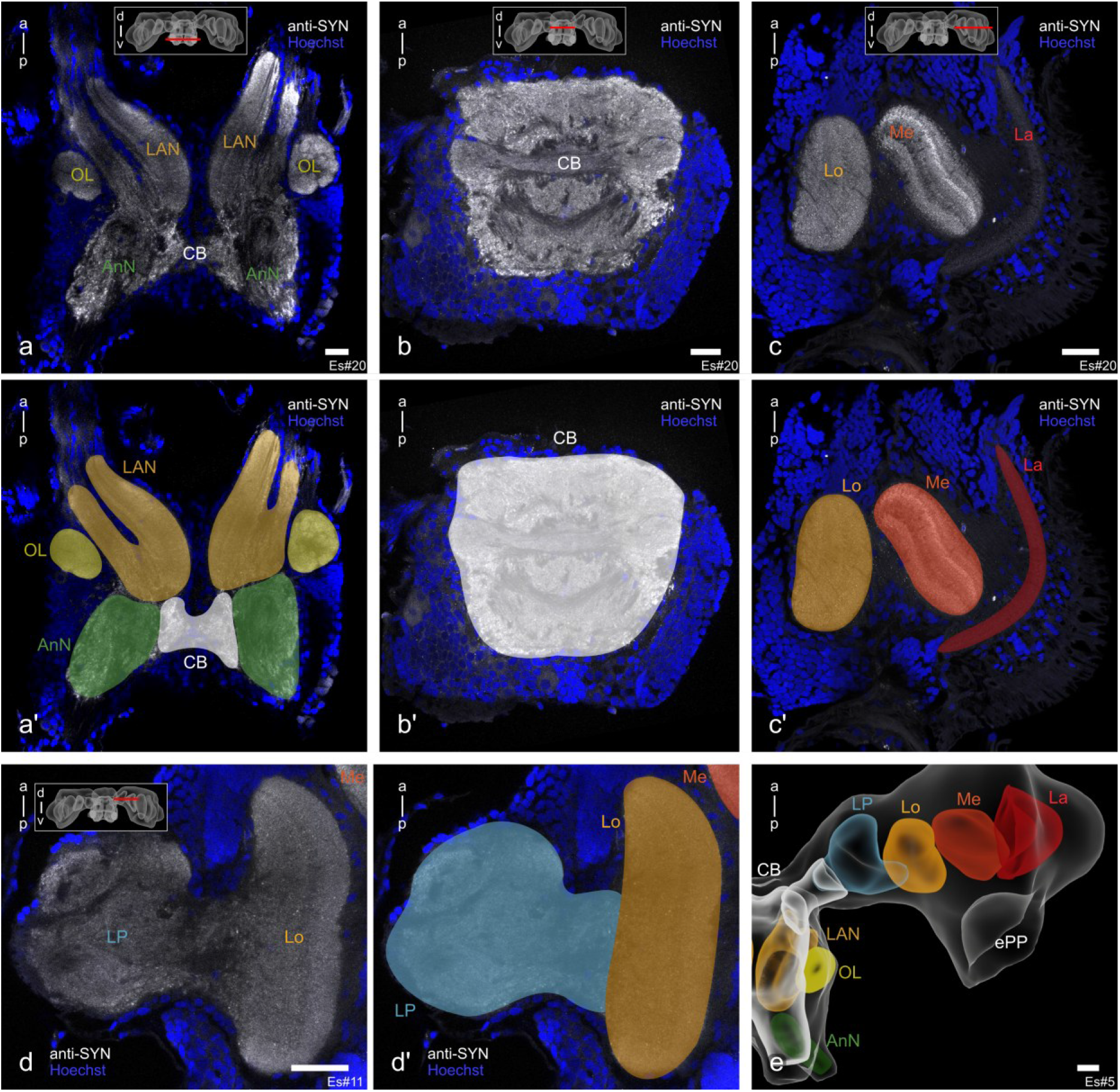
Horizontal vibratome sections of the brain (right eyestalk and central brain) stained against Synapsin (grey) with a nuclear counterstaining (Hoechst, blue) highlight major brain regions. a, a’) Most ventrally, the olfactory neuropils with the lateral antenna 1 neuropil (LAN, ochre), the olfactory lobes (OL, yellow), and the antenna 2 neuropil (AnN, green) are located. **b, b’)** Further dorsally to the olfactory neuropils the central brain (CB) is found. **c-d’)** The eyestalks contain, from lateral to medial, the lamina (La, red), medulla (Me, dark orange), the lobula (Lo, light orange), and the lateral protocerebrum (LP, light blue). **e)** Reconstructions of the neuropils shown for better orientation. The colour scheme is based on Kenning et al. (2013). Scale bars represent 100 µm. Abbreviations: a, anterior; p, posterior; d, dorsal; v, ventral. Figure panels showing the same individual are indicated with the same Es#ID.

**Figure 2-figure supplement 2:**
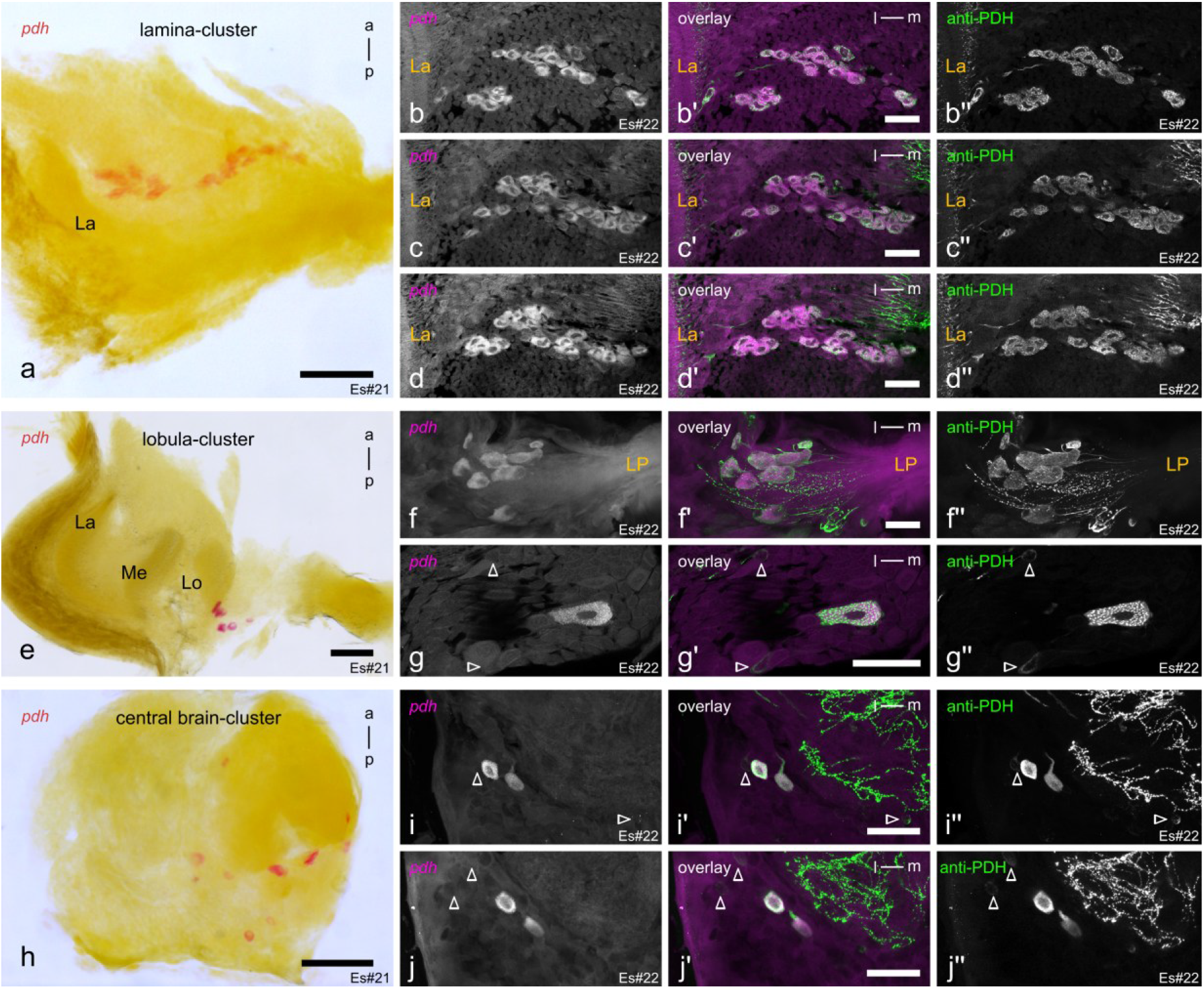
*pdh in situ* hybridization co-localizes with PDH-immunoreactivity. **a)** *pdh in situ* hybridization (red) on a horizontal vibratome section labels a big cell cluster below the lamina (La). **b-d)** single focal planes through the *pdh*-expressing lamina-cell cluster (magenta) from anterior (b) to posterior (d), double labelled with anti-PDH (green), show PDH-immunoreactive (PDH-ir) cells are indeed all expressing *pdh*. **e)** A second cluster of *pdh*-expressing cells is found in the eyestalk, closely associated with the lobula (Lo). **f, g)** Double labelling with anti-PDH shows co-localization of strongly PDH-ir cells and *pdh*-expressing cells also in this lobula-cluster. Occasionally, we found weak PDH-ir cells (blank arrowheads in (g)) where we could not unambiguously identify co-localization between PDH-ir and *pdh*-expression. **h)** *pdh* in situ hybridization reveals several *pdh*-expressing cells in the central brain. **i, j)** Double labelling with anti-PDH shows, like for the other clusters, that strongly immunoreactive cells are also expressing *pdh*. While we could only reliably identify two cells per hemisphere ((i), left; (j), right hemisphere) labelled by in situ hybridization using fluorescence microscopy, we found, for the chromogenic labelling, multiple cells that we might not have been able to distinguish from the strong autofluorescent background in fluorescent microscopy (compare (h, i,j)). Hence, we could not show co-localization for weak PDH-ir cells in the central brain (blank arrowheads in (i, j)), similar to the weakly immunoreactive cells close to the lobula. Scale bars in (a, e, h) represent 250 µm and 100 µm for the other panels. Abbreviations: Me, medulla, LP, lateral protocerebrum; a, anterior; p, posterior; l, lateral; m, medial. Figure panels showing the same individual are indicated with the same Es#ID.

**Figure 2-figure supplement 3:** **Video showing the 3D projection of the entire PDH-ir network.**

**Figure 2-figure supplement 4:** **Video showing single focal planes of the projection of the entire PDH-ir network.**

**Figure 3-figure supplement 1.**
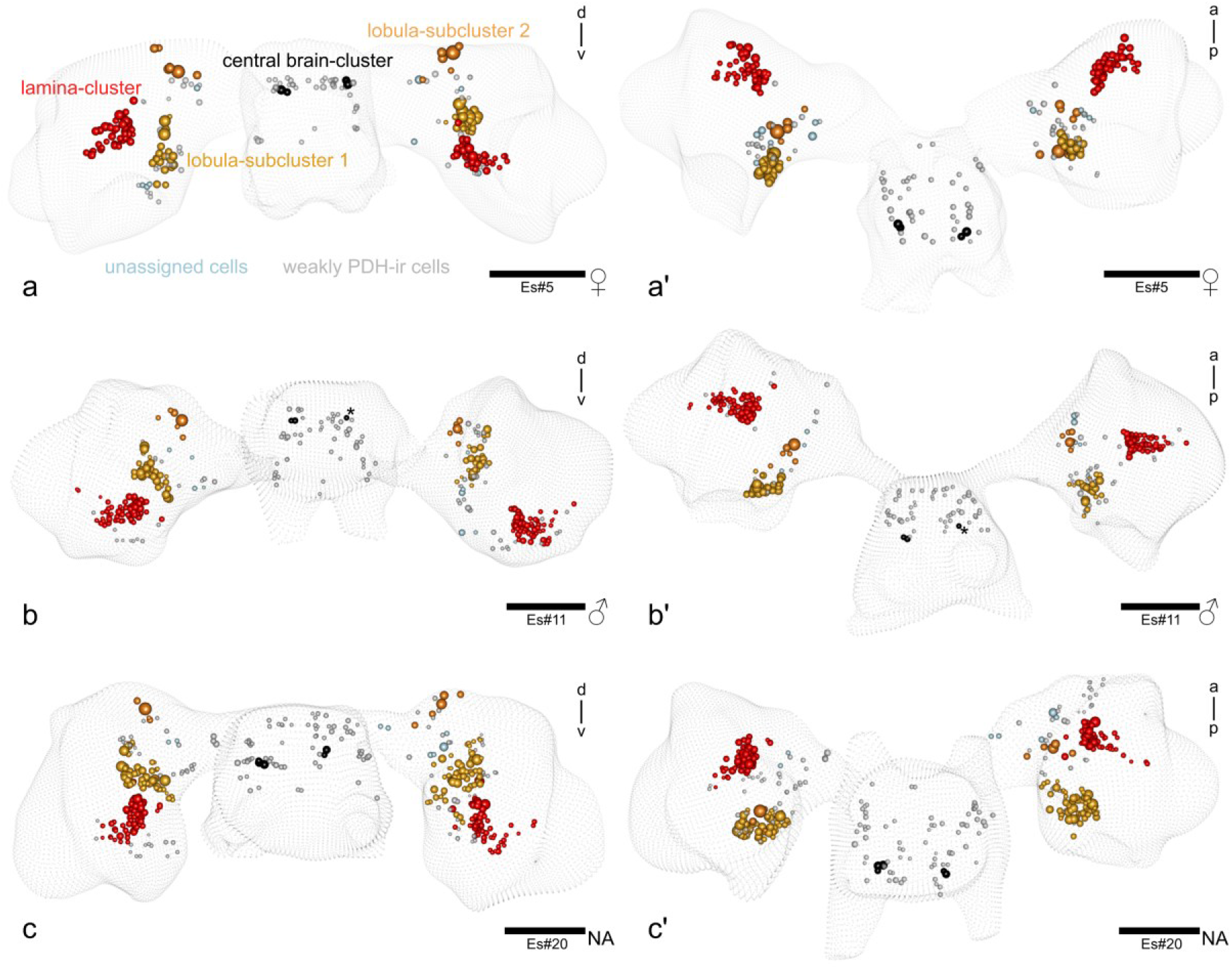
: PDH-ir cell clusters in the brain of krill. a-c) Position of PDH-ir cells within three different brains. Left side always shows the posterior view of the brain (a-c) and right side shows always the dorsal view of the brain (a’-c’). Cells of the lamina-cluster are shown in red, cells of the lobula-subcluster 1 are shown in yellow, cells of lobula-subcluster 2 are shown in orange and strongly PDH-ir cells of the central brain are shown in black. Strongly PDH-ir cells that could not be assigned clearly to any cluster are shown in light blue. Weakly PDH-ir cells that could not be unambiguously shown to co-localize with *pdh* RNA *in situ* hybridization are shown in light grey and are not further considered in the analysis of this paper. Note that the asterisk in (b and b’) indicates a cell of the central brain-cluster that was removed during dissection but its presence could be confirmed by the presence of the distinct projections. Scale bars represent 500 µm. Abbreviations: d, dorsal; v, ventral; a, anterior; p, posterior. Figure panels showing the same individual are indicated with the same Es#ID

**Figure 3-figure supplement 2:** 3D model of PDH-ir cell clusters in the brain of krill. Cells of the lamina-cluster are shown in red, cells of the lobula-subcluster 1 are shown in yellow, cells of lobula-subcluster 2 are shown in orange and strongly PDH-ir cells of the central brain are shown in black. Strongly PDH-ir cells that could not be assigned clearly to any cluster are shown in light blue. Weakly PDH-ir cells that could not be unambiguously shown to co-localize with *pdh* RNA *in situ* hybridization are shown in light grey and are not further considered in the analysis of this paper. Neuropils from Figure 2 as reference. Data for Es#5.

**Figure 4-figure supplement 1:** **Video showing the 3D projection of the PDH-ir network in the central brain.**

**Figure 5-figure supplement 1:**
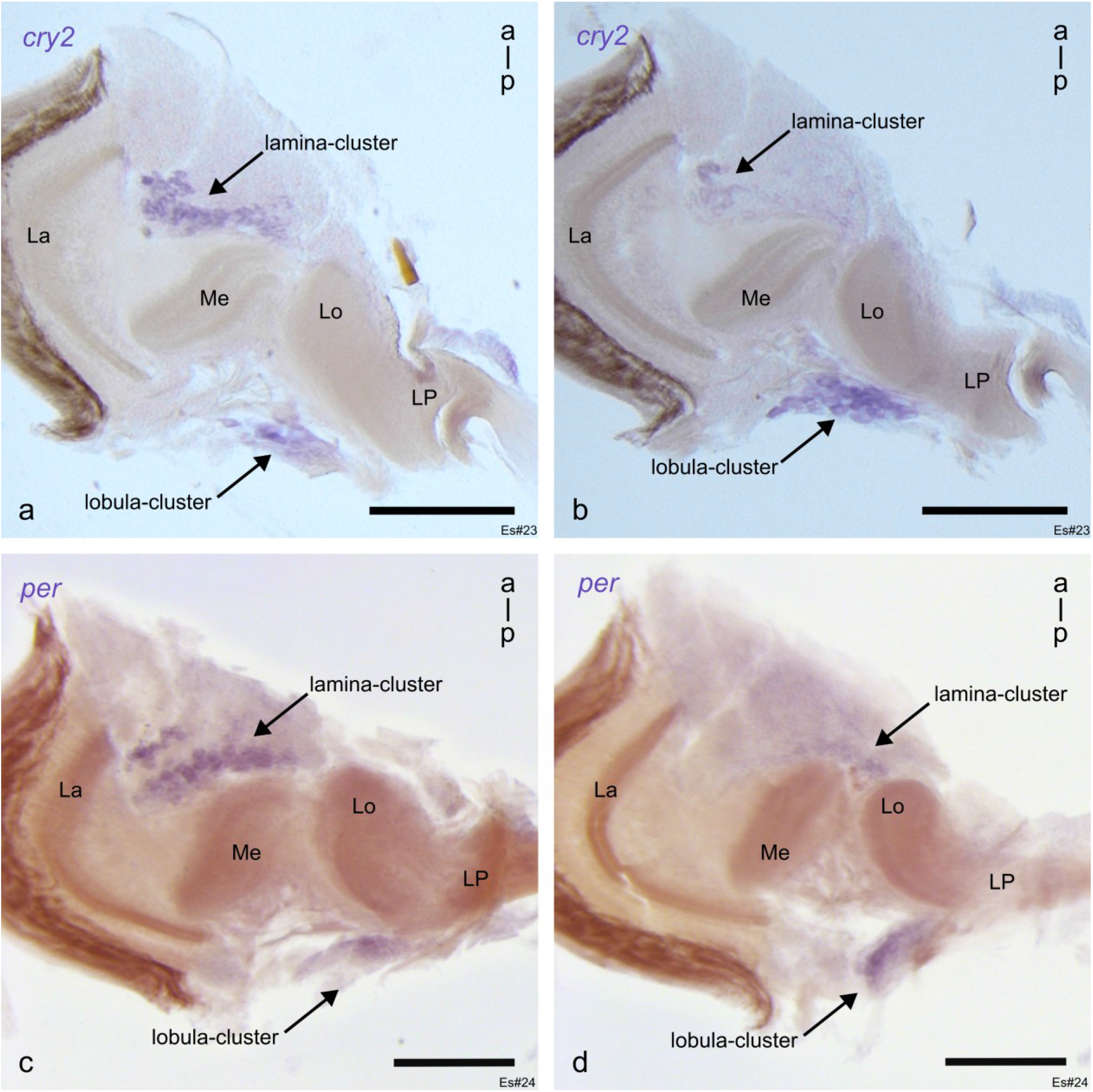
Chromogenic RNA *in situ* hybridization of clock gene mRNA on horizontal vibratome sections of the eyestalk. Bright field image of the *cry2*- **(a, b)** and *per-*expression **(c, d)** in the eyestalks of krill, visualized by chromogenic *in situ* hybridization (violet, arrows). Expression of *cry2* and *per* matches the location of the PDH-positive lamina- and lobula-cluster identified by antibody staining. Scale bars represent 250 µm. Abbreviations: La, lamina; Me, medulla; Lo, lobula; LP, lateral protocerebrum; a, anterior; p, posterior. Figure panels showing the same individual are indicated with the same Es#ID.

**Figure 5-figure supplement 2:**
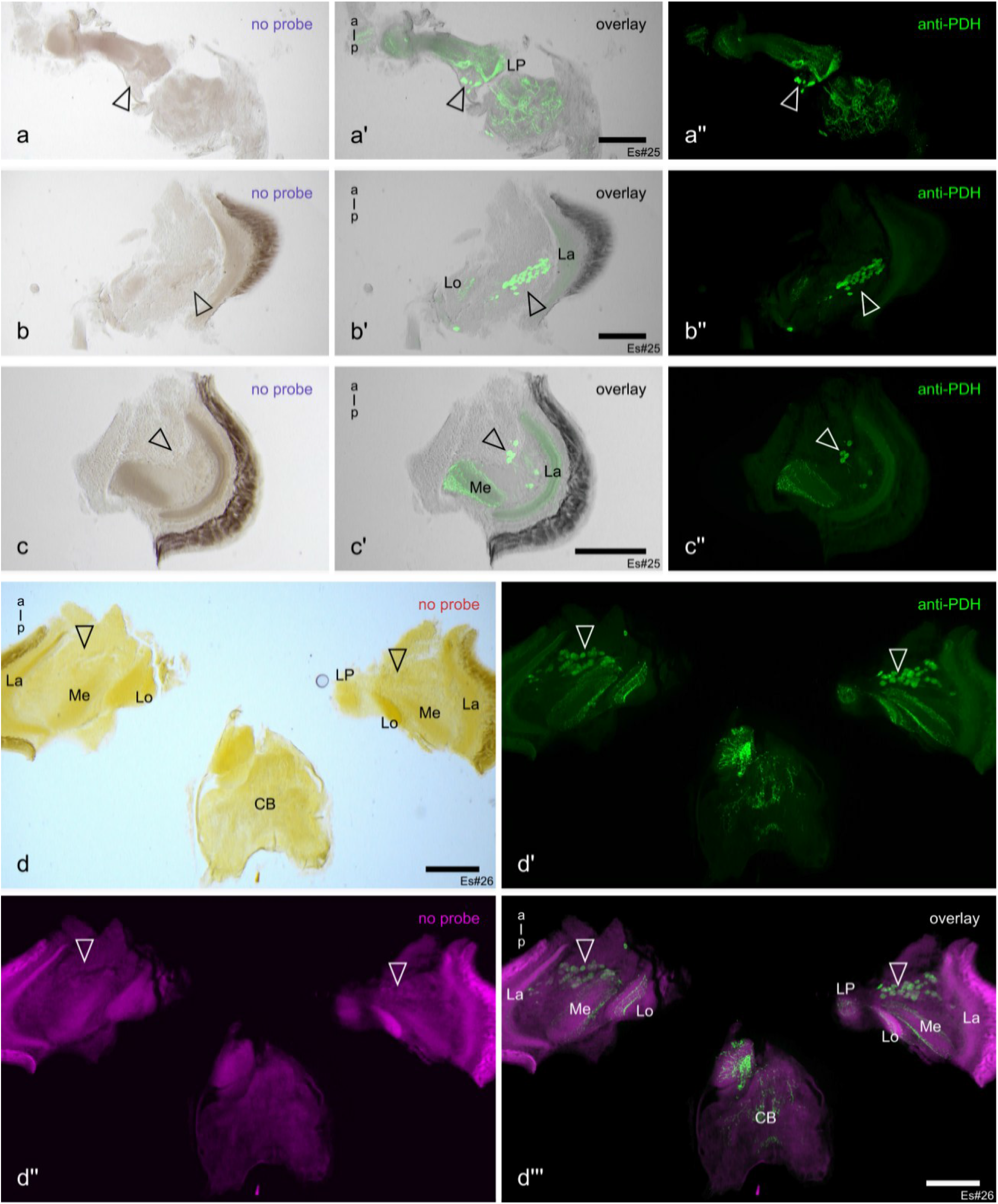
Negative controls for RNA *in situ* hybridization. a-d’’’) Horizontal vibratome sections of krill eyestalks were used in chromogenic *in situ* RNA hybridization without RNA probes as a control for unspecific antibody binding or discoloration of the tissue. **(a-c’’)** sections using BCIP/NBT (violet, (a, b, c)) co-labelled with anti-PDH (green, (a’’, b’’, c’’)) do not show any *in situ* hybridization signal, hence also no co-localization with PDH-positive cells (blank arrowheads, (a’, b’, c’)). This was true for the lobula-cluster (a-a’’) and for the lamina-cluster (b-c’’). **d-d’’’)** Vibratome sections of krill eyestalks used in chromogenic in situ RNA hybridization without RNA probes, stained by Fast Red (red/ magenta, (d, d’’)) and subsequent immunolabeling with anti-PDH (green, (d’, d’’’)), showed, similar to BCIP/NBT, no signal for the *in situ* hybridization and hence no co-localization with anti-PDH (blank arrowheads, d’’’). **d)** brightfield micrograph. **d’-d’’’)** fluorescent micrographs. Scale bars represent 250 µm. Abbreviations: La, lamina; Me, medulla; Lo, lobula; LP, lateral protocerebrum; a, anterior; p, posterior, CB, central brain. Figure panels showing the same individual are indicated with the same Es#ID.

## Supplementary Material

**Supplementary Table 1:**
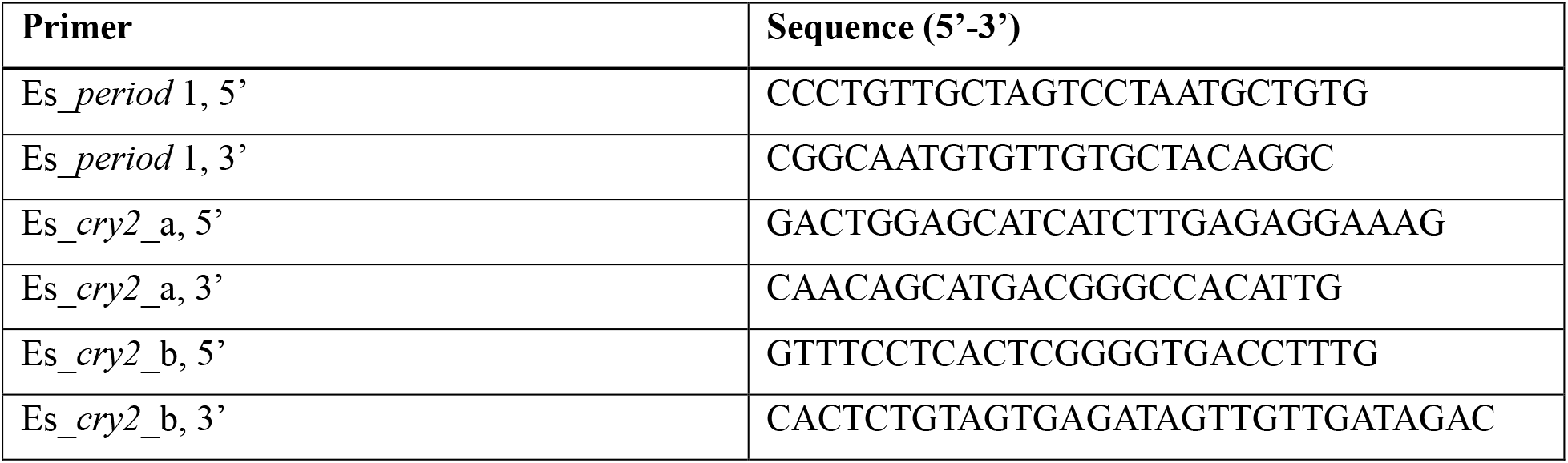
Primer sequences used to synthesize *in situ* hybridization probes. For *cry2* in situ hybridizations, a mix of two probes, termed Es_*cry2*_a and Es_*cry2*_b, was used.

**Supplementary Table 2:** Metadata for each krill Es#ID shown.

| **Krill ID** | **Sex** | **Local Sampling Date** | **Local Sampling Time (UTC-2)** | **Local Sunrise (UTC-2)** | **Local Sunset (UTC-2)** |
|---|---|---|---|---|---|
| **Es#1** | NA | NA | NA | NA | NA |
| **Es#2** | NA | NA | NA | NA | NA |
| **Es#3** | female | 20.03.2022 | 06:10 | 07:07 | 19:18 |
| **Es#4** | female | 20.03.2022 | 03:04 | 07:07 | 19:18 |
| **Es#5** | female | 08/09.02.2022 | 15:00-00:13 | 05:22 | 21:12 |
| **Es#6** | female | 20.03.2022 | 00:07 | 07:07 | 19:18 |
| **Es#7** | NA | 17.03.2022 | 12:10 | 06:57 | 19:27 |
| **Es#8** | male | 02.03.2022 | 03:05 | 06:19 | 20:12 |
| **Es#9** | female | 08/09.02.2022 | 15:00-00:13 | 05:22 | 21:12 |
| **Es#10** | female | 08/09.02.2022 | 15:00-00:13 | 05:22 | 21:12 |
| **Es#11** | male | 28.02./01.03.2022 | 17:50 | 06:13 | 20:18 |
| **Es#12** | male | 09.02.2022 | 00:13 | 05:22 | 21:12 |
| **Es#13** | male | 28.02./01.03.2022 | 14:30-17:50 | 06:13 | 20:18 |
| **Es#14** | male | 28.02./01.03.2022 | 17:50 | 06:13 | 20:18 |
| **Es#15** | male | 21.03.2022 | 05:55 | 07:06 | 19:15 |
| **Es#16** | male | 21.03.2022 | 18:00 | 07:06 | 19:15 |
| **Es#17** | female | 21.03.2022 | 08:45 | 07:06 | 19:15 |
| **Es#18** | male | 21.03.2022 | 18:00 | 07:06 | 19:15 |
| **Es#19** | female | 21.03.2022 | 09:20 | 07:06 | 19:15 |
| **Es#20** | NA | 10.02.2022 | 12:15 | 05:22 | 21:12 |
| **Es#21** | male | 21.03.2022 | 05:55 | 07:06 | 19:15 |
| **Es#22** | female | 20.03.2022 | 06:10 | 07:07 | 19:18 |
| **Es#23** | female | 21.03.2022 | 08:45 | 07:06 | 19:15 |
| **Es#24** | female | 21.03.2022 | 18:00 | 07:06 | 19:15 |
| **Es#26** | female | 21.03.2022 | 09:20 | 07:06 | 19:15 |
| **Es#27** | female | 21.03.2022 | 08:45 | 07:06 | 19:15 |

